# Engineering auxin degradation into root-associated bacteria promotes plant growth

**DOI:** 10.1101/2025.10.25.684584

**Authors:** Ting Jiang, Yihui Shen, Xi Li, Yu Zhou, Michal J. Kozlowski, Philip D. Jeffrey, John T. Groves, Joshua D. Rabinowitz, Jonathan M. Conway

## Abstract

Overproduction of indole-3-acetic acid (IAA) by rhizosphere bacteria disrupts plant auxin homeostasis and induces root growth inhibition (RGI). *Variovorax* reverses this effect by degrading IAA, but the underlying pathway remains incompletely resolved. Here, using genetics, metabolomics, and isotope tracing, we define a nine-gene region (*iadCDEFGHIJK2*) required for IAA catabolism in *Variovorax paradoxus* CL014, identify a previously uncharacterized intermediate (C₁₀H₉NO₃), and revise the early oxidative steps of the pathway. Although IadDE adopts a Rieske-type oxygenase architecture, our data are consistent with the IadCDE complex functioning as a monooxygenase during IAA degradation. Using these insights, we introduce *iad* genes into two rhizobacterial chassis, *Polaromonas* MF047 and *Paraburkholderia* MF376. These engineered strains degrade IAA and alleviate RGI induced by exogenous IAA, an auxin-producing strain, and a synthetic bacterial community. Engineered *Paraburkholderia* MF376 delivers the strongest performance, improving plant growth in natural soil. Together, these results establish a framework for engineering auxin-balancing root commensals.

## Introduction

Plant roots develop in a densely populated microbial environment in which soil and rhizosphere microbes strongly influence root architecture and function^1,2^. A central regulator of root development is auxin, a group of signaling molecules that function as plant hormones but are also synthesized by bacteria, fungi, and animals^3^. Indole-3-acetic acid (IAA) is the predominant naturally occurring auxin^4^. Auxin responses are highly dose- and tissue-dependent^5–7^: low concentrations promote growth, whereas higher concentrations inhibit it. This biphasic response is especially pronounced in roots, which are more sensitive than shoots and display growth promotion only within a narrow concentration range; increases above this optimum can trigger root growth inhibition (RGI)^7^. Plants maintain auxin homeostasis through coordinated control of auxin biosynthesis, transport, conjugation, and degradation^8^. However, IAA produced by rhizosphere bacteria can perturb this balance and thereby inhibit root growth^9^. Genomic surveys indicate that more than 80% of soil- and plant-associated bacterial genomes encode complete or partial IAA biosynthetic pathways^10,11^, highlighting bacterially derived auxin as a widespread and important component of plant–microbe interactions in soil.

Certain rhizosphere bacteria can mitigate RGI by degrading IAA. Two major types of auxin-degrading pathways have been identified in soil- or plant-associated bacteria: the *iad*-like and *iac*-like pathways^12,13^. The *iad*-like (IAA degradation) pathway, found in genera such as *Variovorax*, *Alcaligenes*, *Achromobacter,* and *Bradyrhizobium*, converts IAA into anthranilic acid^9,12^. *Variovorax* is broadly distributed across root-associated microbial communities, and the *iad*-like locus is highly conserved within the genus^9,12,14^. In contrast, the *iac*-like (indole-3-acetic acid catabolism) pathway, originally described in *Pseudomonas putida* 1290, degrades IAA into catechol^15,16^. Compared with the *iac*-like pathway, the *iad*-like pathway is more effective in reversing RGI induced by root-associated bacteria^12^. For example, *Variovorax* strains, including *V. paradoxus* CL014, carrying the *iad* pathway fully restored root growth in *Arabidopsis thaliana* seedlings exposed to a 175-member RGI-inducing synthetic community, whereas *iac*-containing strains did not^12^. This protective effect was also observed in sorghum, where addition of *Variovorax* to an RGI-inducing synthetic community improved plant performance in a soil-based drought phenotyping experiment^17^.

Despite its critical role, the molecular mechanisms underlying *iad*-mediated IAA degradation remain incomplete. The *iad* locus is known to be regulated by two MarR-family transcription factors, MarR73 and MarR50, with MarR73 serving as the primary repressor^12^. The first catalytic step of the pathway involves IadDE, annotated as a Rieske non-heme dioxygenase, and the associated reductase IadC. Together, IadCDE are postulated to form a two-component dioxygenase system that facilitates electron transfer and enhances catalytic efficiency^12,18^.

Previous studies have demonstrated the feasibility of engineering IAA-degrading activity by introducing *iad* genes into heterologous hosts^9,12,18^. Plasmid-based expression of the *iad* locus in the root-associated isolate *Acidovorax* Root219 conferred IAA-degrading activity *in vitro* and promoted primary root elongation under exogenous IAA treatment^9^. However, this *Acidovorax* Root219 strain failed to fully reverse RGI caused by the auxin-producing *Arthrobacter* CL028, let alone more complex auxin-producing bacterial communities^9^. These findings highlight the need to identify more suitable chassis strains for *iad*-pathway engineering, particularly strains that are better adapted to complex microbial communities and can exhibit enhanced growth-promoting functions during plant–microbe interactions when equipped with *iad* genes.

Here, we investigated the genetic and metabolic basis of *iad*-mediated IAA degradation in *V. paradoxus* CL014 and evaluated the transferability of this pathway into additional root-associated bacterial chassis. By combining genetics, metabolomics, and isotope tracing, we defined the core genes required for IAA catabolism, identified new pathway intermediates, and revised the previously proposed *iad* degradation pathway at the metabolite level. Using this information, we identified and engineered a chassis strain that, when equipped with the *iad* genes, effectively performs auxin-balancing functions. This engineered strain is better adapted to colonization and persistence in complex rhizosphere microbial communities, enabling plant growth promotion in contexts where *V. paradoxus* CL014 is less effective.

## Results

### Identification of a nine-gene region within the *iad* locus responsible for IAA degradation in *Variovorax paradoxus* CL014

The *iad* locus (Hot Spot 33, HS33), a 25-gene region, was linked to IAA degradation in CL014 by genomic deletion (Fig. 1a)^9^. To pinpoint key genes, we cloned different segments of the *iad* locus into the broad-host vector pBBR1 and introduced them into an *iad* locus deletion mutant (*V. paradoxus* CL014 ΔHS33) (Extended Data Fig. 1a). Metabolomic analysis revealed that mutants carrying the *iadCDEFGHIJK2* (*iadC-K2*) region restored the level of intermediates and the final product, anthranilic acid (C₇H₇NO₂), to that observed in the wild type, indicating that this nine-gene region encodes the complete IAA degradation pathway (Fig. 1a, Extended Data Fig. 1b).

**Fig. 1.**
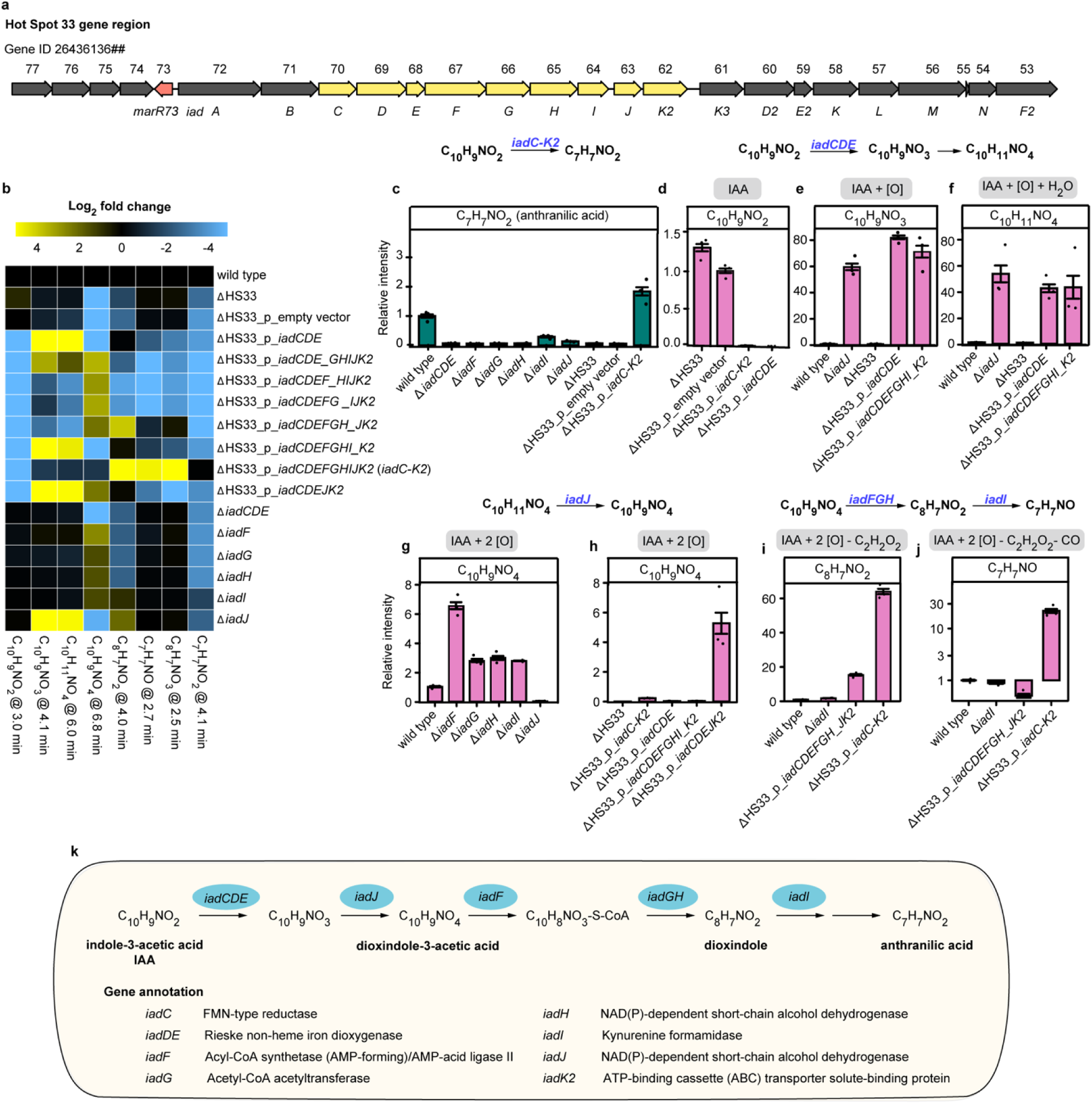
Metabolomic profiling of *iad* pathway mutants reveals substrates and products of genes involved in IAA degradation in *V. paradoxus* CL014. **a,** Genomic organization of Hot Spot 33 in *V. paradoxus* CL014. Gene annotations are shown above as the final two digits of the IMG gene ID (26436136##), with assigned *iad* gene names shown below. The MarR-family transcriptional regulator *marR73* is shown in red; the core nine-gene IAA degradation locus is highlighted in yellow. **b**, Heatmap of Log₂ fold changes in metabolite abundance in *iad* pathway mutants relative to the wild type. Full LC–MS profiles are provided in Extended Data Fig. 2. Data represent the mean of *n* = 3 biological replicates. **c–j**, Bar plots showing the relative abundance of anthranilic acid (**c**), IAA (**d**), and key metabolic intermediates (**e–j**) across wild-type and mutant strains. Data represent mean ± s.e.m. (standard error of the mean) of *n* = 3 biological replicates. **k**, Schematic representation of the *iad*-mediated IAA degradation pathway in *V. paradoxus* CL014.

To identify specific substrates and products of each Iad enzyme, we performed liquid chromatography in tandem with high-resolution mass spectrometry (LC-MS), focusing on metabolites previously associated with IAA degradation in *V. paradoxus* CL014^12^. We reasoned that knocking out a gene downstream of a metabolite or overexpressing a gene upstream would lead to metabolite accumulation or depletion, respectively. Accordingly, we created knockout mutants in the wild-type background and overexpression mutants in the ΔHS33 background, and measured metabolite levels (Fig. 1b-j, Extended Data Fig. 2). Based on gene annotations and structural data, *iadD* and *iadE* encode the large and small subunits of a Rieske non-heme dioxygenase, while *iadC* encodes a reductase, forming a two-component dioxygenase system (Fig. 1k)^12,18^. Accordingly, *iadCDE* was treated as a single functional unit for mutant construction.

Our analysis revealed that deletion of *iadCDE*, *iadF*, *iadG*, *iadH*, *iadI*, or *iadJ* significantly reduced anthranilic acid (C₇H₇NO₂) production (Fig. 1c), confirming their essential roles in IAA degradation. Complementation of the ΔHS33 mutant with the full *iadC–K2* gene region fully restored anthranilic acid levels (Fig. 1c). Notably, the presence of *iadCDE* in these segments was able to rapidly degrade IAA. This confirms that IadCDE directly uses IAA as its substrate and catalyzes the initial step of IAA degradation (Fig. 1d, Extended Data Fig. 1c). We next examined metabolite accumulation in the ΔHS33_p_*iadCDE* (p, plasmid-borne expression) strain and identified two compounds, C₁₀H₉NO₃ and likely its hydration product C₁₀H₁₁NO₄, suggesting C₁₀H₉NO₃ to be the product of IadCDE (Fig. 1e,f). Interestingly, *iadJ* deficiency also led to the accumulation of C₁₀H₉NO₃ and C₁₀H₁₁NO₄, indicating that IadJ likely consumes one of these intermediates (Fig. 1e). Similarly, we found that C₁₀H₉NO₄ is depleted with *iadJ* deletion while it accumulates in ΔHS33_p_*iadCDEJK2*, suggesting C₁₀H₉NO₄ to be the product of IadJ (Fig. 1g,h). Deletion of *iadF*, *iadG*, *iadH* and *iadI* all led to C₁₀H₉NO₄ accumulation, among which Δ*iadF* shows the strongest accumulation suggesting functional proximity of IadF to IadCDEJ (Fig. 1g). Other projected intermediates were not detected, thus IadFGHI likely carry out a chain of reactions in which most intermediates are rapidly channeled through the enzymes. Therefore, we resorted to annotated enzyme function to infer reactions carried out by these gene products. Annotated as an acyl-CoA synthase (AMP-forming), IadF likely activates the carboxylic group in the acetyl side chain, enabling its subsequent removal by IadG, an acetyl-CoA acetyltransferase (ketothiolase, EC 2.3.1.16) (Fig. 1k). This reaction yields CoA-bound intermediates that are membrane-impermeable, ensuring their retention within bacterial cells and enabling efficient recognition and processing^19^. Meanwhile, IadH, annotated as an alcohol dehydrogenase, likely facilitates further processing (Fig. 1k). Together, IadF, IadG, and IadH mediate the reductive removal of the acetyl side chain (-C₂H₂O₂), producing C₈H₇NO₂ (Fig. 1i). Indeed, C₈H₇NO₂ accumulates in the ΔHS33_p_*iadCDEFGH*_*JK2* expression strain as well as the *iadI* deletion strain (Fig. 1i), suggesting that IadI acts downstream of C₈H₇NO₂. We also detected C₇H₇NO and C₈H₇NO₃, which strongly accumulate only in strains overexpressing the whole *iadC-K2* gene locus, suggesting that they are likely intermediates between the IadI product and the final product anthranilic acid (C₇H₇NO₂) (Fig. 1b, j).

Although *iadI* is annotated as a kynurenine formamidase, our MS/MS analysis confirmed that C₈H₇NO₃ is not N-formylanthranilic acid—the expected formylated derivative of anthranilic acid—indicating that IadI may instead function as a broad-activity amidase (Extended Data Fig. 1d). Lastly, IadK2 was previously identified as a highly IAA-specific ATP-binding cassette (ABC) transporter solute-binding protein involved in IAA uptake^18^. However, it is not essential, as ΔHS33_p_*iadCDE* was still capable of taking up and utilizing IAA, suggesting alternative uptake mechanisms^12^.

In summary, we identified a nine-gene region (*iadC-K2*) within the *iad* locus responsible for IAA degradation in *V. paradoxus* CL014 (Fig. 1a, Extended Data Fig. 1a,b). By integrating metabolite abundance changes with gene functional annotations, we assigned putative reaction steps to each gene and reconstructed the degradation pathway (Fig. 1k). This pathway links specific functional genes to corresponding intermediates, providing a framework for understanding IAA metabolism in this microorganism.

### New intermediate in two-step oxidation for IAA degradation

Previous models proposed that *iad*-mediated IAA degradation proceeds through monooxygenation of IAA to form 2-oxindole-3-acetic acid (oxIAA, C₁₀H₉NO₃), followed by conversion to dioxindole^12,18^. However, our metabolomic analysis revealed a new uncharacterized intermediate, rather than oxIAA, to be the product of monooxygenation (Extended Data Fig. 3). Specifically, we detected two distinct C₁₀H₉NO₃ chromatogram peaks in wild-type extracts: a major peak at 4 min and a minor peak at 7 min (Extended Data Fig. 3a). The minor peak was confirmed to be oxIAA by comparison with a chemical standard (Extended Data Fig. 3a); however, its abundance was not reduced in the ΔHS33 mutant, indicating that oxIAA is unlikely to be the product of *iad*-mediated IAA oxidation (Extended Data Fig. 3b,c). On the contrary, the major C₁₀H₉NO₃ peak, hereafter referred to as **compound 1**, was depleted in ΔHS33 and accumulated in *iadCDE*-overexpressing strains, supporting its assignment as the IadCDE-derived product (Extended Data Fig. 3b,c).

To gain mechanistic insight into the oxidation reactions, we performed ^2^H and ^18^O isotope tracing and LC-MS/MS analysis of both labeled and unlabeled pathway intermediates (Fig. 2, Extended Data Fig. 3d-f and 4). Specifically, we first employed uniformly deuterium-labeled IAA ([²H₇]IAA) to monitor hydrogen rearrangements during catabolism. Mass shifts in pathway intermediates and their MS fragments reveal the number and position of retained deuterium atoms, providing greater mechanistic resolution than the previously used carbon-labeled IAA ([¹³C₆]IAA)^12^. The first intermediate, C₁₀H₉NO₃, retained all seven deuterium atoms, suggesting no hydrogen loss during the initial oxidation step (Fig. 2a, Extended Data Fig. 4a,c). This observation is inconsistent with mechanisms involving oxIAA or its enol form, 2-hydroxy-IAA^12,18^, or a radical intermediate from hydrogen atom abstraction^20^. Indeed, the minor C₁₀H₉NO₃ peak corresponding to oxIAA retained only six deuterium atoms (Extended Data Fig. 3d). The subsequent intermediate, C₁₀H₉NO₄, showed the loss of one deuterium, whereas all downstream intermediates consistently retained four deuterium atoms on the aromatic ring (Fig. 2a, Extended Data Fig. 4a,c).

**Fig. 2.**
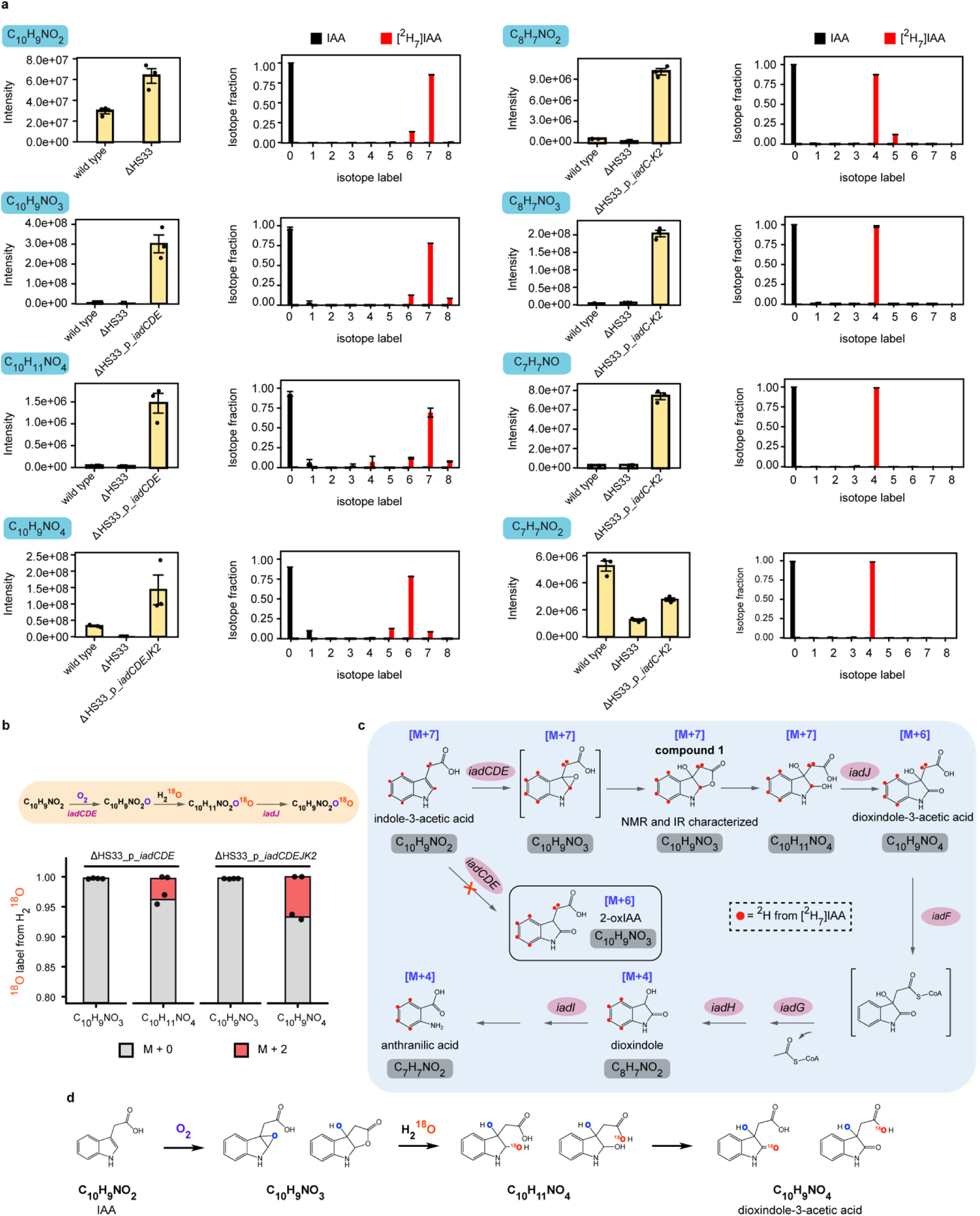
Isotope tracing with [²H₇]IAA and H ^18^O reveals a revised IAA degradation pathway in *V. paradoxus* CL014. **a**, LC-MS analysis of [²H₇]IAA catabolism identifies key pathway intermediates and their deuterium labeling profiles. Left, LC–MS peak intensities of major metabolites. Right, deuterium isotope distributions following incubation with unlabeled IAA or [²H₇]IAA. Data represent mean ± s.e.m. (standard error of the mean) of *n* = 3 biological replicates. **b,** H₂¹⁸O (20% v/v) tracing confirms the source of oxygen atoms incorporated during the two-step oxidation reactions. Data are shown for two biological replicates. **c**, A revised mechanistic model of IAA degradation consistent with isotope tracing. Structures in brackets are hypothesized. **d**, Proposed chemical structures of two-step oxidation products and their transformations, consistent with observed mass shifts and isotope retention patterns.

We next sought to identify the source of oxygen in the formation of C₁₀H₉NO₃, C₁₀H₁₁NO₄, and C₁₀H₉NO₄ using ¹⁸O-labeled water (H₂¹⁸O) tracing in bacterial culture (Fig. 2b, Extended Data Fig. 4b). ^18^O tracing distinguishes oxygenase mechanisms (O from molecular oxygen) and dehydrogenase mechanisms (O from water). No ^18^O labeling was detected in **compound 1** (C₁₀H₉NO₃) in the *iadCDE*-overexpressing strain, consistent with IadCDE catalyzing monooxygenation during formation of this intermediate (Fig. 2b). By contrast, oxIAA was significantly labeled by H₂¹⁸O, supporting that oxIAA is not formed by an oxygenase (Extended Data Fig. 3e). The subsequent products, C₁₀H₁₁NO₄ and C₁₀H₉NO₄, also incorporated ¹⁸O, supporting a model in which C₁₀H₁₁NO₄ is formed via hydration of C₁₀H₉NO₃ in cells, and is further processed by IadJ to C₁₀H₉NO₄ through a dehydrogenase mechanism (Fig. 2b). Further MS/MS analysis of C₁₀H₉NO₄ revealed that the incorporated ¹⁸O resides either on the carboxylate or the C2 ketone moiety of the dioxindole-3-acetic acid (Fig. 2d, Extended Data Fig. 4b,c). Interestingly, C₁₀H₉NO₄ remained almost unlabeled from ^18^O tracing in the wild type (Extended Data Fig. 3f), which may be due to the rapid hydrolysis of C₁₀H₉NO₃ within the IadCDE catalytic center, where H₂O is derived from catalytic reduction of molecular oxygen possibly by IadC.

From the data above, we hypothesized that the product of IadCDE, **compound 1** (C₁₀H₉NO₃), is a rearranged lactone following epoxidation of IAA (Fig. 2c)^21^. This mechanism and the proposed lactone product agree with full preservation of seven deuterium atoms in C₁₀H₉NO₃ from [²H₇]IAA, as well as the H_2_^18^O tracing pattern in C₁₀H₉NO₄ (Fig. 2d). To validate the hypothesized structure, we chemically synthesized **compound 1** and validated its structure by nuclear magnetic resonance spectroscopy (NMR) and infrared spectroscopy (IR) (Extended Data Fig. 3g and Supplementary Information). We then confirmed using LC-MS/MS that the intracellular intermediate indeed structurally matched with **compound 1** (Extended Data Fig. 3h) in both retention time and MS/MS fragmentation. Moreover, incubation of the synthesized **compound 1** with ΔHS33_p_*iadCDEJK2* cell lysate resulted in formation of the downstream product dioxindole-3-acetic acid (C₁₀H₉NO₄) (Extended Data Fig. 3i).

Together, these findings indicate that the *iad* pathway in *V. paradoxus* CL014 proceeds through a two-step oxidation process. The first oxidation, achieved by *iadCDE*, converts IAA to a previously uncharacterized monooxygenated intermediate, **compound 1**, likely through a 2,3-epoxide mechanism^21,22^ followed by intramolecular rearrangement leading to lactone formation. The subsequent oxidation is achieved by IadJ through a dehydrogenase mechanism (Fig. 2c).

### IadCDE adopts a Rieske-type architecture and is sufficient for IAA degradation and RGI reversal

Metabolomic and isotope-tracing analyses indicated that IadCDE functions as a monooxygenase complex. To investigate the structure of this enzyme, we solved the crystal structure of the IadDE complex from CL014 at 1.28 Å resolution (PDB: 9O71; Fig. 3a), revealing the canonical architecture of a Rieske non-heme oxygenase complex. Phylogenetic analysis further placed IadD, the catalytic subunit, in a distinct subclade within the broader Rieske dioxygenase and indole oxygenase family^23^. IadD is most closely related to the phthalate family of Rieske non-heme oxygenases, but also shows relationships to members of the heme-dependent aromatic oxygenase (HDAO) and flavin-dependent monooxygenase (FMO) superfamilies (Extended Data Fig. 5), suggesting shared functional features across distinct oxygenase lineages.

**Fig. 3.**
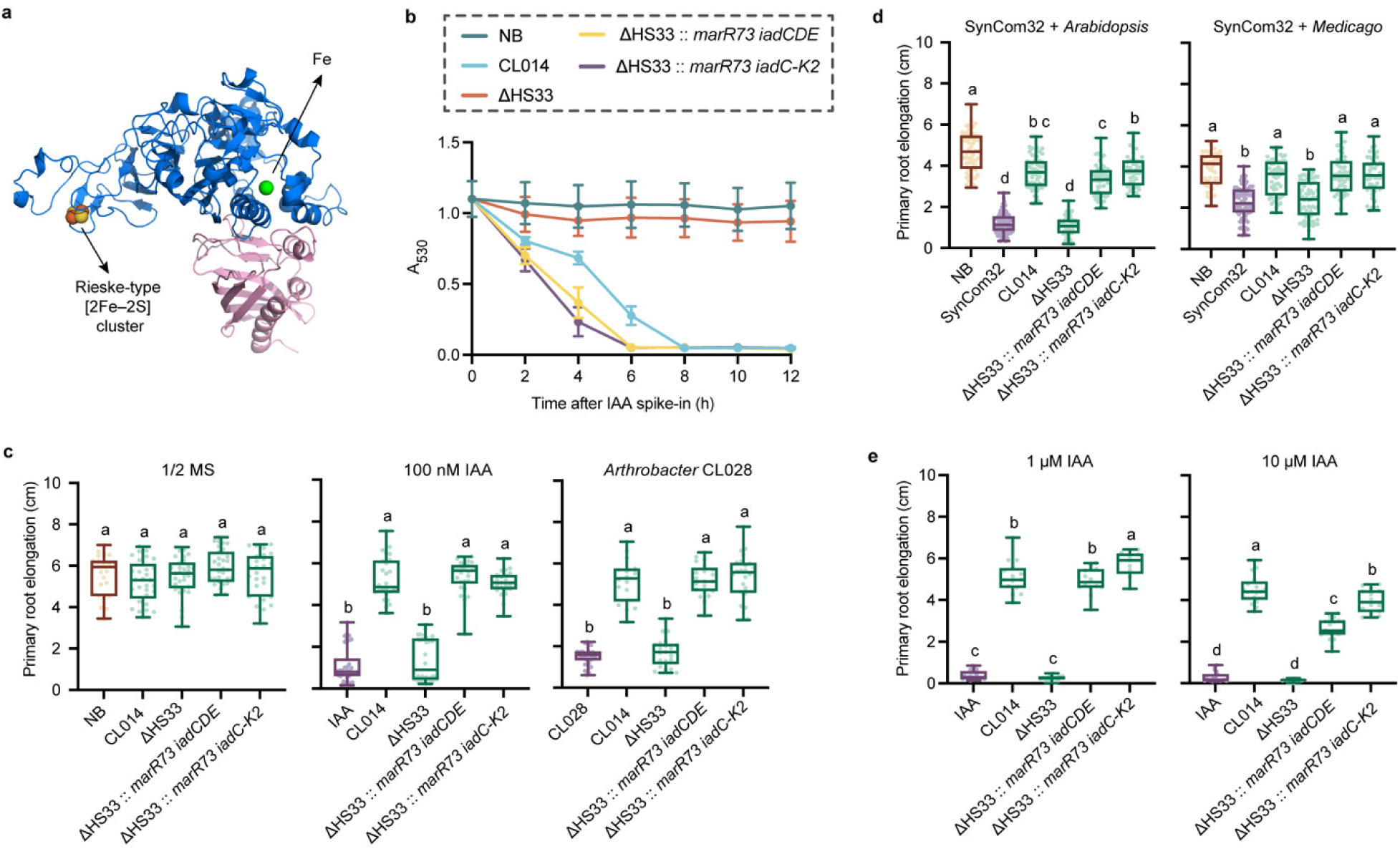
Structural and genetic evidence show that IadCDE enables auxin degradation and alleviates root growth inhibition. **a**, Crystal structure of the IadDE complex from *V. paradoxus* CL014 at 1.28 Å resolution (PDB code: 9O71), with IadD shown in blue and IadE in pink. **b**, *in vitro* IAA degradation by engineered strains. IAA concentrations were measured from culture supernatants of strains grown in M9 minimal medium supplemented with glucose and 0.1 mg/mL IAA. The Salkowski reagent reacts with IAA to generate a pink-to-red chromophore with an absorbance maximum at 530 nm (A₅₃₀). Data represent the mean of two biological replicates per sample. **c–e**, Primary root length of *Arabidopsis* seedlings exposed to 100 nM IAA or the auxin-producing strain *Arthrobacter* CL028 (**c**), A 32-member synthetic community (SynCom32) in *Arabidopsis* and *Medicago* seedlings (**d**), and elevated IAA concentrations (1 μM and 10 μM) in *Arabidopsis* (**e**). Letters above box plots indicate statistically significant differences determined by one-way ANOVA with Tukey’s post hoc test on the underlying replicate values (*P* < 0.05). Groups that do not share a letter are significantly different. “NB” denotes the no-bacteria control. Sample sizes (left to right): **c**, *n* = 23, 29, 31, 31, 27, 34, 26, 34, 26, 25, 31, 16, 21, 21, 25; **d**, *n* = 64, 81, 56, 52, 53, 43, 52, 78, 52, 71, 47, 48; **e**, *n* = 19, 15, 18, 14, 9, 19, 17, 19, 19, 14. Box plots represent the median (center line), interquartile range (box), and 1.5 × interquartile range (whiskers).

A homologous structure (PDB: 7YLS) was previously resolved at 1.8 Å resolution, with 98.4% and 100% sequence identity with CL014 IadD and IadE, respectively^18^. Comparison of the two structures revealed two main local differences. First, residue 280 of IadD is histidine in our structure but alanine in the previously reported model, and this difference is associated with an altered loop orientation between the structures. Second, the loop spanning residues 234–240 of IadD also adopts a different orientation. Both regions map to the active site^18^, and differences in this architecture may influence enzymatic activity. Thus, our higher-resolution structure refines local active-site structural features rather than redefining the overall fold of IadDE.

Because oxygenases are central to bacterial utilization of aromatic compounds, we next asked whether the IadCDE module alone is sufficient to support IAA degradation and mitigate RGI. To test this, we genomically integrated the native regulator *marR73*^12^ together with either *iadCDE* or the full pathway (*iadC-K2*) into the ΔHS33 background. This design enabled direct comparison of the oxygenase module with the complete degradation pathway. In bacterial culture medium supplemented with 0.1 mg ml⁻¹ (570 μM) IAA, both engineered strains showed substantially enhanced IAA degradation relative to the wild type, with the full-pathway strain showing slightly faster degradation than the *iadCDE* strain (Fig. 3b). In a gnotobiotic plant system, however, both strains fully alleviated RGI to a similar extent in *Arabidopsis* challenged with 100 nM exogenous IAA, the auxin-producing strain *Arthrobacter* CL028^12^, or a 32-member bacterial synthetic community (SynCom32; Supplementary Table 1), and comparable effects were observed in *Medicago* (Fig. 3c,d). These results show that the IadCDE module alone is sufficient to reverse RGI under these conditions.

However, under higher IAA concentrations in the plant assay (1 μM or 10 μM), the strain carrying the full *iadC-K2* pathway outperformed the minimal *iadCDE* strain. In addition, strains carrying only *iadCDE* showed slightly reduced growth relative to strains carrying the full pathway in bacterial culture supplemented with 0.1 mg ml⁻¹ (570 μM) IAA, indicating that downstream pathway steps contribute to more robust detoxification under high-IAA conditions (Fig. 3e and Extended Data Fig. 6e).

### Chromosomal integration of *iad* genes enables IAA degradation and RGI reversal in root-associated bacteria

To evaluate whether root-associated bacteria can be engineered to degrade IAA, we selected two root-associated strains for *iad* pathway integration: *Polaromonas* MF047 and *Paraburkholderia* MF376^24^. Both strains, like *Variovorax*, are Betaproteobacteria and do not impair host growth when co-inoculated with *Arabidopsis* seedlings (Fig. 4b). Neither strain produced IAA when cultured in M9 minimal medium supplemented with tryptophan, nor were they able to reverse RGI triggered by exogenous IAA or by the auxin-producing strain *Arthrobacter* CL028 (Fig. 4b, Extended Data Fig. 6a). Additionally, *Polaromonas* MF047 exhibited amino acid-dependent growth, showing no growth in M9 minimal medium unless supplemented with specific amino acids (Extended Data Fig. 6b).

**Fig. 4.**
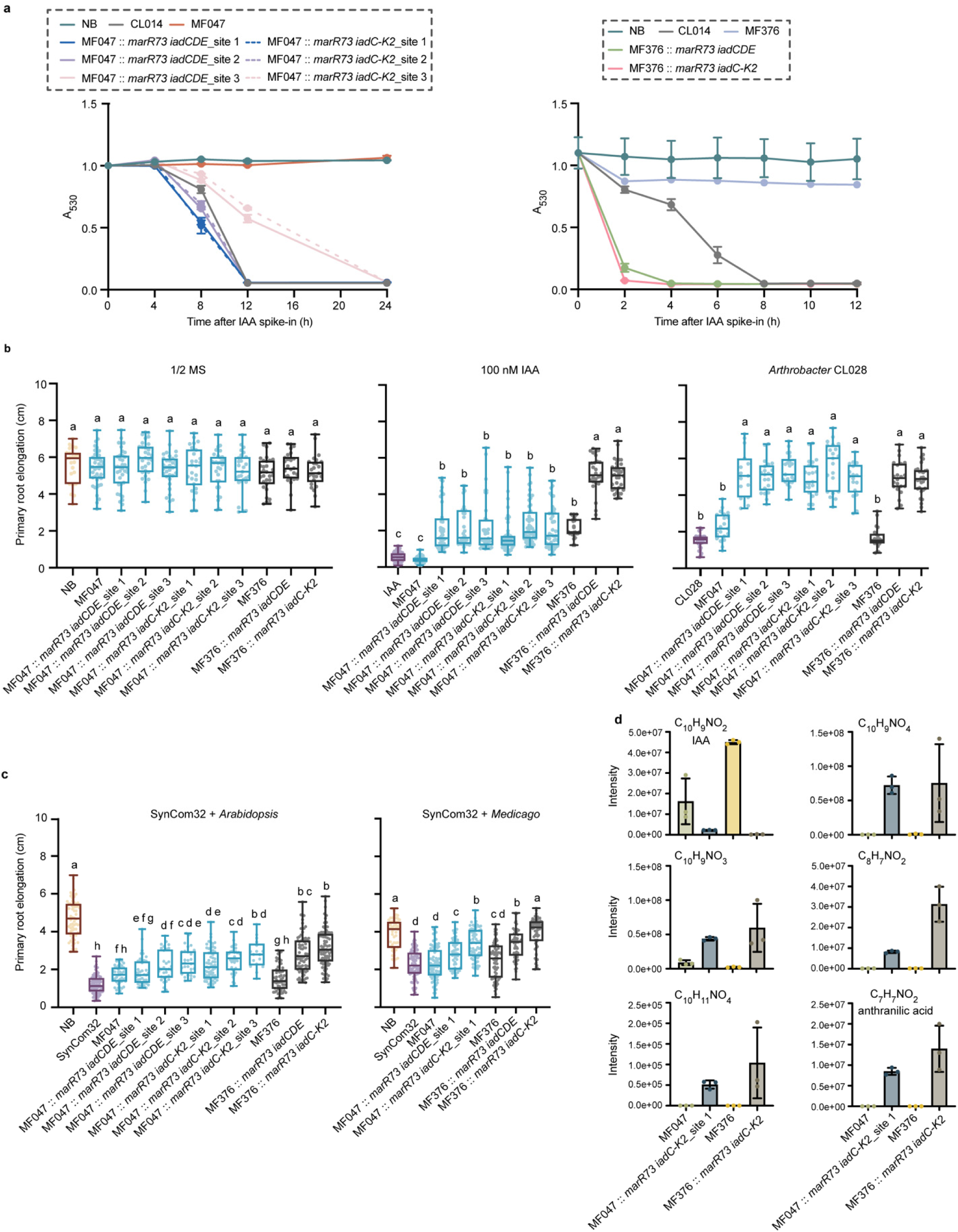
Engineering root-associated bacteria with *iad* genes enables IAA degradation and alleviates root growth inhibition (RGI). **a**, Engineered *Polaromonas* MF047 strains cultured in 50% TSB medium and *Paraburkholderia* MF376 strains cultured in M9 medium efficiently degraded IAA. IAA levels in culture supernatants were quantified using the Salkowski reagent, which reacts with IAA to form a pink-to-red chromophore with an absorbance maximum at 530 nm (A₅₃₀). Data represent the mean of two biological replicates per sample. **b–c**, Engineered chromosomal knock-in strains reversed RGI, as indicated by enhanced primary root elongation in *Arabidopsis* seedlings treated with 100 nM IAA or the auxin-producing strain *Arthrobacter* CL028 (**b**), and in *Arabidopsis* and *Medicago* seedlings exposed to a 32-member synthetic bacterial community (SynCom32) (**c**). Letters above box plots indicate statistically significant differences determined by one-way ANOVA with Tukey’s post hoc test on the underlying replicate values (*P* < 0.05). Groups that do not share a letter are significantly different. “NB” denotes the no-bacteria control. Sample sizes (left to right): **b**, *n* = 23, 42, 37, 31, 34, 26, 30, 27, 31, 26, 25, 51, 42, 44, 33, 39, 52, 49, 42, 26, 28, 35, 31, 21, 15, 21, 21, 23, 21, 17, 22, 22, 29; **c**, *n* = 64, 81, 45, 41, 33, 28, 61, 36, 17, 56, 75, 89, 52, 78, 76, 58, 67, 74, 57, 68. Box plots show the median (horizontal line), interquartile range (box), and whiskers extending to 1.5 × the interquartile range. **d**, Bar plots of the LC-MS peak intensities of IAA, pathway intermediates, and the final product (anthranilic acid) of the *iad* pathway in MF047 (50% TSB) and MF376 (M9 medium) and their corresponding strains carrying the full-pathway knock-in construct (*marR73 iadC-K2*) (*n* = 3).

To confer IAA-degrading capacity, we genomically integrated *marR73* together with either the minimal (*iadCDE*) or complete (*iadC-K2*) pathway into three intergenic sites in MF047 and one site in MF376 (Extended Data Fig. 6c). To minimize potential disruption of native gene expression, insertion sites were located within non-coding intergenic regions ranging from 400 to 900 bp in length. All engineered strains degraded IAA within 24 hours, and MF376 knock-ins achieved complete IAA removal within four hours (Fig. 4a). This rapid degradation is consistent with the observation that RGI was alleviated in *Arabidopsis* seedlings exposed to 100 nM exogenous IAA (Fig. 4b). Growth analysis in 50% TSB medium with or without IAA showed no significant differences between wild-type and engineered strains, indicating that chromosomal integration did not impair bacterial fitness under the tested conditions (Extended Data Fig. 6d,f,g).

To assess their function in plant-microbe interactions, we co-inoculated engineered strains with *Arthrobacter* CL028 or SynCom32 on 7-day-old *Arabidopsis* seedlings in a gnotobiotic system. After 7 days, all engineered strains fully rescued RGI triggered by *Arthrobacter* CL028, restoring primary root length to control levels (Fig. 4b). In parallel, they significantly alleviated RGI induced by SynCom32, with MF376 consistently showing greater efficacy than MF047 (Fig. 4c). To test whether this activity extends across host species, we repeated the SynCom32 experiment using *Medicago*. Under these conditions, strains carrying the full *iad* pathway promoted greater primary root elongation than strains expressing only the minimal *iadCDE* module (Fig. 4c), highlighting the importance of complete IAA degradation for robust root growth promotion in complex microbial environments.

To determine whether the engineered strains degrade IAA through the same pathway as CL014, we profiled pathway-associated metabolites by LC–MS and isotope tracing. In both strains carrying the full *iad* pathway, IAA was depleted relative to the corresponding wild type, and the expected intermediates as well as the final product anthranilic acid (C₇H₇NO₂) accumulated and were detected, supporting the conclusion that the engineered strains degrade IAA through the same pathway as CL014 at the metabolomic level (Fig. 4d). However, unlike in CL014, where anthranilic acid can be further used for tryptophan biosynthesis (Extended Data Fig. 6h), we did not detect evidence that *iad*-derived anthranilic acid is incorporated into tryptophan in the engineered strains (Extended Data Fig. 6i). Although these strains are able to synthesize tryptophan, this synthesis does not appear to proceed through the *iad*-derived anthranilic acid (Extended Data Fig. 6i).

Together, these results show that stable genomic integration of *iad* genes into root-associated bacteria is sufficient to confer IAA degradation and restore root growth under microbial auxin stress. The engineered strains function across distinct microbial contexts and in multiple plant hosts, supporting their utility as tools to modulate plant-associated auxin metabolism.

### Engineered *Paraburkholderia* MF376 reverses RGI that *V. paradoxus* CL014 cannot fully rescue

To assess the contribution of the *iad* pathway to RGI mitigation, we selected 23 previously identified RGI-inducing strains from nine bacterial genera (Supplementary Table 1)^9^. Each strain was co-inoculated with either wild-type *V. paradoxus* CL014 or the *iad* deletion mutant ΔHS33 on 7-day-old *Arabidopsis* seedlings. After 7 days, root length measurements revealed that 22 strains induced RGI (Fig. 5a, b; Extended Data Fig. 7). Among these, RGI caused by 16 strains was reversed by wild-type *V. paradoxus* CL014, and in 12 of these cases (75%), the wild type restored root growth more effectively than the *iad*-deficient mutant ΔHS33, indicating an *iad*-dependent mitigation mechanism (Extended Data Fig. 7). In contrast, for *Agrobacterium* MF224, *Arthrobacter* MF135, and *Pseudomonas* MF051, both wild-type and ΔHS33 strains restored root growth to similar levels, suggesting an *iad*-independent mechanism (Fig. 5b).

**Fig. 5.**
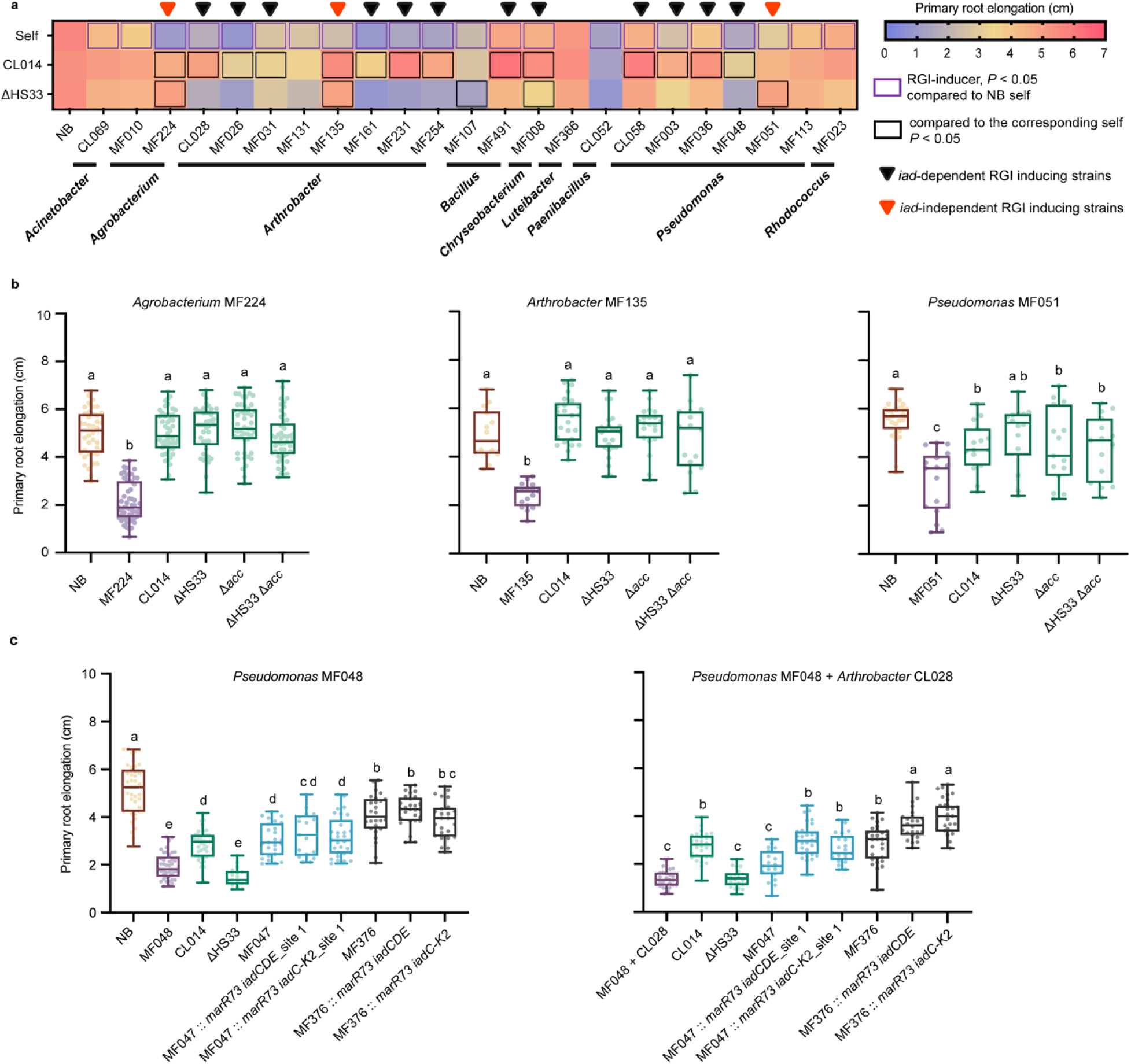
Root growth inhibition by certain strains is mitigated independently of the *iad* locus, while engineered *Paraburkholderia* MF376 enhances mitigation of *iad*-dependent inhibition. **a,** Heatmap showing average primary root lengths of *Arabidopsis* seedlings inoculated with 23 previously identified RGI-inducing strains^9^, either alone (self) or co-inoculated with *V. paradoxus* CL014 wild type or the *iad*-deficient mutant ΔHS33, to assess *iad*-dependent reversion. Blue squares indicate significant differences compared to the no-bacteria (NB) control demonstrating RGI phenotypes; black squares indicate significant differences compared to the strain inoculated alone (self). Corresponding box plots are shown in Extended Data Fig. 7. **b**, Primary root length of *Arabidopsis* seedlings co-inoculated with *iad*-independent RGI-inducing strains and treated with *V. paradoxus* CL014 wild type, ΔHS33, Δ*acc* (ACC deaminase gene deleted), or ΔHS33 Δ*acc* (both *iad* and ACC deaminase genes deleted). **c**, Primary root lengths of *Arabidopsis* seedlings inoculated with *Pseudomonas* MF048 alone or with *Arthrobacter* CL028, and co-inoculated with *V. paradoxus* CL014, or *iad*-engineered strains of *Polaromonas* MF047 or *Paraburkholderia* MF376. Letters above box plots indicate statistically significant differences determined by one-way ANOVA with Tukey’s post hoc test on the underlying replicate values (*P* < 0.05). Groups that do not share a letter are significantly different. “NB” denotes the no-bacteria control. Sample sizes (left to right): **b**, *n* = 40, 56, 48, 41, 48, 49, 14, 17, 25, 25, 21, 16, 28, 17, 15, 14, 15, 15; **c**, *n* = 42, 41, 41, 17, 25, 17, 29, 27, 29, 25, 26, 28, 22, 19, 30, 21, 30, 24, 25. Center lines, boxes, and whiskers indicate the median, interquartile range, and 1.5 × interquartile range, respectively.

Genome analysis identified a gene in CL014 encoding ACC deaminase, which degrades ACC, the precursor of ethylene^25^. Because ethylene can inhibit primary root elongation at high levels^26^, and prior work showed that RGI induced by *Arthrobacter* CL028 and a synthetic microbial community requires both auxin and ethylene signaling in the host^9^, we tested whether ACC deaminase contributes to the *iad*-independent rescue phenotype. However, deletion of the ACC deaminase gene in both wild-type and ΔHS33 backgrounds did not impair RGI alleviation, ruling out ACC deaminase as the primary mechanism underlying this *iad*-independent effect (Fig. 5b). These results indicate that although the *iad* pathway is a major mechanism by which CL014 suppresses RGI caused by many root-associated strains, CL014 also uses additional *iad*-independent mechanisms against a subset of microbes.

Despite its broad effectiveness, CL014 was unable to fully rescue RGI induced by *Pseudomonas* MF048. To test whether *iad*-engineered strains could provide enhanced reversion, we evaluated MF047 and MF376 engineered with the *iad* pathway. These strains were co-inoculated with *Pseudomonas* MF048 alone or in combination with *Arthrobacter* CL028 on *Arabidopsis* seedlings. Root length analysis showed that MF376 knock-in strains significantly outperformed both CL014 and engineered MF047 under both conditions (Fig. 5c). Collectively, these results demonstrate that engineering *Paraburkholderia* MF376 with the *iad* pathway enhances its ability to mitigate complex RGI interactions, supporting its potential as a promising chassis bacterium for plant growth promotion.

### Impact of *V. paradoxus* CL014 and engineered strains on the root microbiome and plant growth

CL014 can degrade IAA without harming plant growth at normal treatment levels (OD₆₀₀ = 0.05). To test whether this effect holds under high bacterial load, we applied CL014 at OD₆₀₀ = 2 to *Arabidopsis* seedlings in a gnotobiotic system. No significant changes in primary root length were observed, indicating that even at high densities, CL014 does not adversely affect plant growth or disturb auxin balance (Fig. 6a). These findings suggest that its IAA-degrading function may primarily buffer auxin produced by neighboring microbes rather than directly altering plant root growth.

**Fig. 6.**
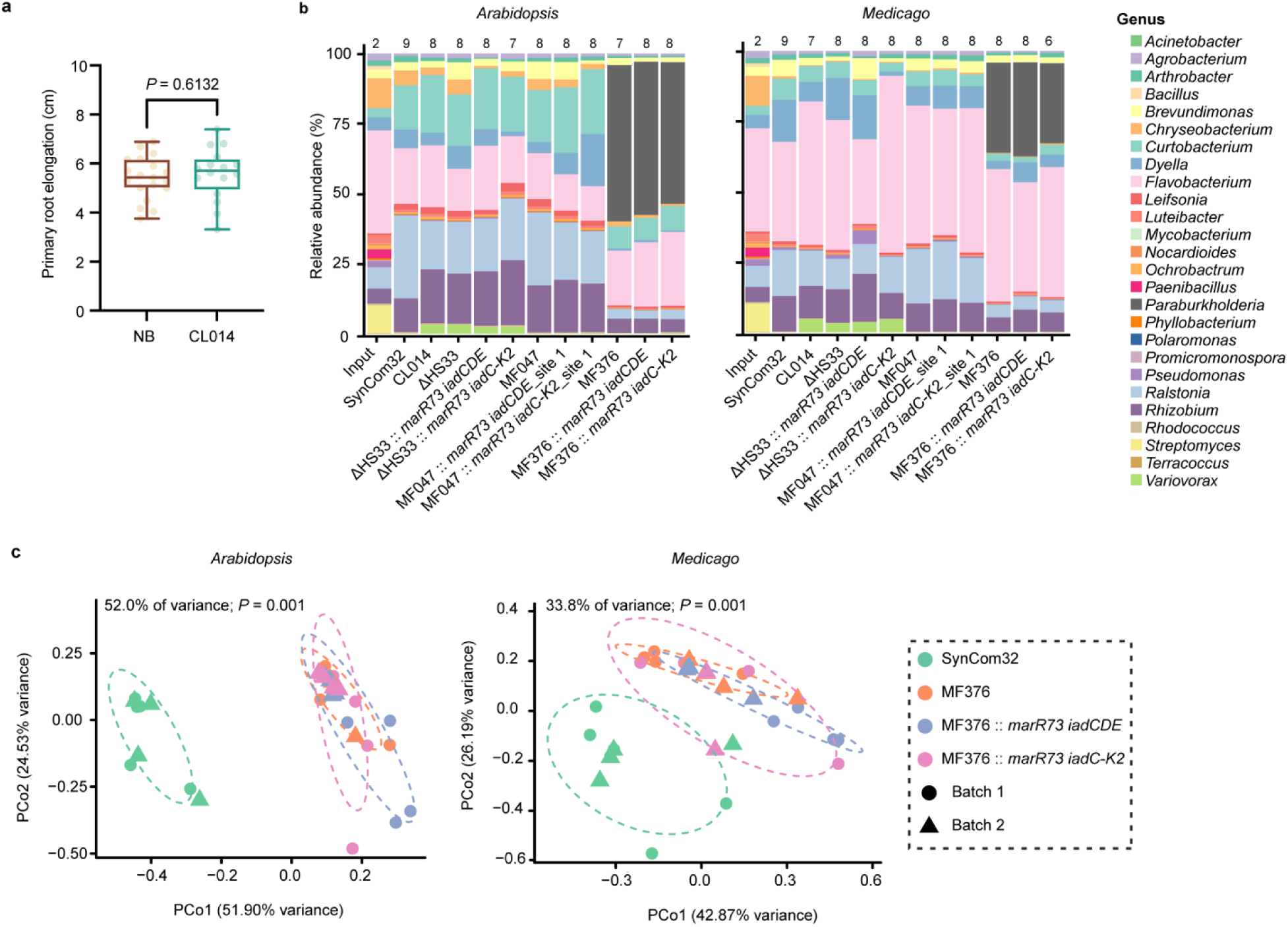
The bacterial *iad* pathway does not significantly alter overall SynCom composition under the tested conditions. **a**, Primary root length of *Arabidopsis* seedlings treated with *V. paradoxus* CL014 at OD₆₀₀ = 2. Statistical significance was assessed using Welch’s two-tailed *t*-test (*n* = 19, 16). **b**, Relative abundance of bacterial genera in the root microbiomes of *Arabidopsis* and *Medicago* seedlings treated with SynCom32 alone or supplemented with wild-type or *iad*-engineered strains of *V. paradoxus* CL014, *Polaromonas* MF047, and *Paraburkholderia* MF376. Sample sizes are indicated above each bar. **c**, Unconstrained principal coordinate analysis (PCoA) of Bray–Curtis dissimilarity showing root microbiome profiles in *Arabidopsis* and *Medicago* seedlings treated with SynCom32 alone or with engineered *Paraburkholderia* MF376 strains. Ellipses represent 68% confidence intervals. Statistical significance was determined by PERMANOVA (Adonis2). Data shown in panels **a–c** are derived from two independent experiments.

To assess the impact of CL014, MF047, MF376, and their *iad*-gene-containing engineered strains on the root microbiome, we treated *Arabidopsis* and *Medicago* seedlings with SynCom32 (Supplementary Table 1) alone or supplemented individually with each wild-type or *iad*-engineered strain. After 9 days, root-associated communities were profiled by 16S rRNA amplicon sequencing. Despite receiving the same SynCom32 inoculum, *Arabidopsis* and *Medicago* developed distinct root microbiomes, highlighting the influence of host genotype (Fig. 6b). Among the three introduced genera, *Paraburkholderia* exhibited the highest relative abundance in roots (50.84%), followed by *Variovorax* (3.40%), whereas *Polaromonas* (0.01%) showed minimal colonization (Fig. 6b, and Supplementary Table 4). Due to its robust root colonization, both wild-type and engineered *Paraburkholderia* MF376 substantially altered root microbiome composition, explaining 52% and 33.8% of the community variation between SynCom32 alone and SynCom32 supplemented with MF376-related strains in *Arabidopsis* and *Medicago*, respectively, as shown by unconstrained principal coordinate analysis (PCoA) using Bray–Curtis distances (*P* = 0.001; Fig. 6c). However, no significant or consistent differences in overall community composition were detected between wild type and *iad*-engineered MF376 strains. In contrast, *Variovorax* CL014- and *Polaromonas* MF047-related treatments produced modest shifts in community composition (17.1-28.3% explained variation, *P* < 0.05), consistent with their lower colonization (Fig. 6b, Extended Data Fig. 8 and Supplementary Table 4). Likewise, log₂ fold-change analysis revealed no significant differences in the relative abundance of any bacterial genus between wild-type and engineered derivative strains within the same chassis background (Extended Data Fig. 9). Together, these results show that no significant or consistent differences in overall community composition were detected between wild-type and corresponding iad-engineered strains within the same chassis background. In contrast, larger community-level differences were observed among the different bacterial chassis and were associated with their markedly different relative abundances.

To assess potential benefits for plant performance, we inoculated untreated natural soil with each strain and grew *Arabidopsis* and *Medicago* for 33 days. For *Medicago*, we quantified shoot fresh weight, shoot dry weight, root fresh weight, root dry weight and root length. Because *Arabidopsis* roots were too fragile to be reliably recovered intact from soil, only shoot fresh and dry weight were quantified for this species. Plants grown in soil inoculated with MF376 carrying the complete *iad* pathway (*iadC-K2*) showed significant increases in shoot and root biomass as well as root length compared with uninoculated controls and plants treated with the wild-type strain (Fig. 7 and Extended Data Fig. 10). Although differences between the full-pathway strains and the minimal-cassette strains (*iadCDE*) were not always statistically significant, the full-pathway strains consistently showed a stronger growth-promotion trend. Together, these results indicate that combining a strongly root-colonizing chassis with the complete IAA degradation pathway yields the most robust growth-promotion phenotype in natural soil under the conditions tested here.

**Fig. 7.**
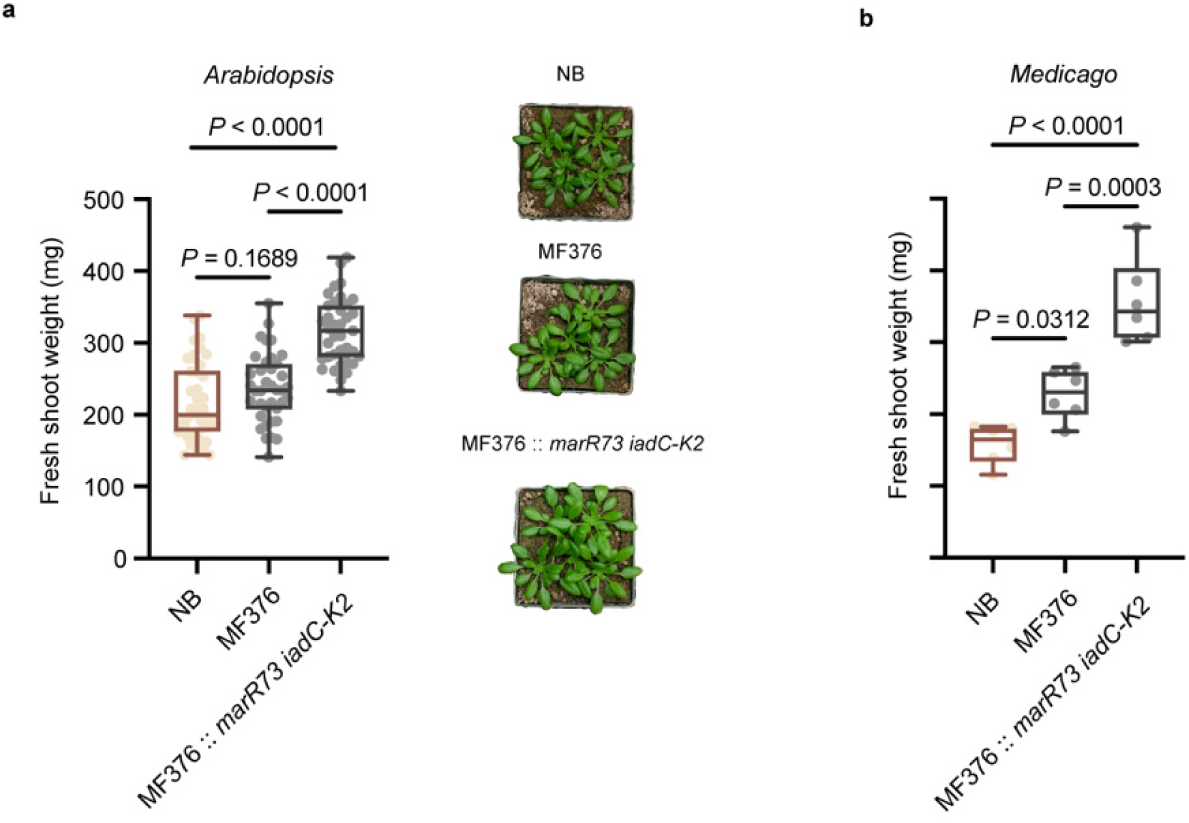
*Paraburkholderia* MF376-derived engineered strains promote plant growth during long-term cultivation in natural soil. Fresh shoot weight of 33-day-old *Arabidopsis* (a) and *Medicago* (b) plants grown in untreated natural soil inoculated individually with *Paraburkholderia* MF376 wild type and *marR73 iadC-K2* knock-in strains. Statistical significance was determined using one-way ANOVA with Tukey’s post hoc test (*n* = 42, 38, 37, 6, 6, 6). Data are derived from four independent experiments for *Arabidopsis* and two independent experiments for *Medicago*.

## Discussion

Members of the genus *Variovorax* have been shown to reverse RGI by degrading auxin produced by root-associated bacteria^9^. In *V. paradoxus* CL014, a 25-gene region, the *iad* locus, was previously identified as being required for this phenotype^9^. However, the roles of individual genes and the sequence of metabolic transformations in the IAA degradation pathway remained incompletely defined. The ability of this pathway to function in more complex plant-microbe contexts had also not been established. Here, using genetics, metabolomics, and isotope tracing, we delineated the *iad*-mediated IAA degradation pathway in CL014 at both the genetic and metabolite levels, identifying new intermediates and revising the previously proposed pathway. Using this understanding, we introduced *iad* genes into two root-associated chassis strains and evaluated their ability to degrade IAA, alleviate auxin-associated RGI, and promote plant growth across diverse contexts. Engineered *Paraburkholderia* MF376 emerged as the more effective chassis across these contexts, including in conditions where CL014 was less effective. This highlights the potential of transferring a mechanistically defined auxin-degradation pathway into alternative root-associated strains to improve plant outcomes.

A central advance of this work is the revision of the early oxidative steps of the *iad* pathway (Fig. 2). Previous studies, based largely on metabolite assignments from [¹³C₆]IAA tracing, proposed that CL014 degrades IAA through a route analogous to that described in *Bradyrhizobium japonicum*, proceeding through 2-hydroxyindole-3-acetic acid, dioxindole-3-acetic acid, isatin, isatinic acid and anthranilic acid^12,27^. Because the major structural changes occur on the pyrrole ring and acetic acid side chain, we instead used [²H₇]IAA to resolve these transformations. This revealed a different early pathway architecture, in which the initial IAA oxidation proceeds through two sequential steps (C₁₀H₉NO₂ → C₁₀H₉NO₃ → C₁₀H₉NO₄) (Fig.1 and 2). IadCDE catalyses the first oxidation using oxygen derived from molecular oxygen, whereas IadJ acts downstream in formation of C₁₀H₉NO₄, with the additional oxygen atom derived from water (Fig. 2). Our data further show that the C₁₀H₉NO₃ intermediate is structurally distinct from the commonly assumed intermediate oxIAA^18^, despite sharing the same molecular formula (Extended Data Fig. 3). These revisions provide a more accurate framework for understanding *iad*-like pathways in plant-associated bacteria from genera such as *Variovorax*, *Bradyrhizobium*, *Alcaligenes* and *Achromobacter*^12^.

Oxygenases play a crucial role in aerobic bacteria by incorporating oxygen into chemically stable aromatic compounds, enabling their degradation into metabolically accessible substrates^19^. To date, the most extensively studied indole oxygenases fall into two major families: the heme-dependent aromatic oxygenase (HDAO) superfamily, which catalyzes dioxygenation or monooxygenation via an Fe/heme-dependent mechanism^22,28,29^; and the flavin-dependent monooxygenases (FMOs), which use a flavin cofactor to mediate oxygen transfer^30–32^. Rieske-type dioxygenases can exhibit diverse catalytic activities, functioning as dioxygenases, monooxygenases, or hydroxylases^20,28,33^. Our metabolomics and isotope tracing reveal a monooxygenation-driven mechanism, identifying IadCDE in CL014 as a two-component indole monooxygenase that is essential for initiating IAA degradation, rather than a dioxygenase as previously annotated^12,18^. The proposed reaction mechanism closely resembles that of MarE, a heme-dependent aromatic monooxygenase involved in the maremycin biosynthetic pathway of *Streptomyces* sp. B9173^22,28,29,34^. While MarE belongs to the HDAO superfamily and relies on a heme cofactor^22,28^, IadDE is structurally similar to Rieske non-heme dioxygenases (Fig. 3a, b)^18^.

IadC and IadDE together form a two-component oxygenase system in which IadC transfers electrons from NADH to IadDE to activate molecular oxygen^18,35^. Although IadC enhances catalytic efficiency^12^, it is not strictly required for detectable IAA degradation in CL014 under simplified liquid culture conditions, likely because endogenous reductases can partially compensate for its function. Consistent with this, overexpression of *iadDE* alone in the ΔHS33 background restored IAA degradation in bacterial culture assays^12^, and IadDE activity was further found to be strain-dependent when expressed heterologously, functioning in *Escherichia coli*10 Beta but not in *Escherichia coli* BL21(DE3), likely owing to host-specific differences in redox environment^12,18^. Similar functional redundancy has been reported in *Sphingopyxis granuli*, where ThnA4, a ferredoxin reductase, is dispensable for tetralin degradation^36^. However, the requirement for IadC differs in the plant context: in *Arabidopsis*, only ΔHS33 strains expressing *iadCDE*, but not *iadDE* alone, were able to reverse RGI induced by the auxin-producing *Arthrobacter* CL028^12^. Together, these results indicate that although IadC can be bypassed for IAA degradation in simplified liquid assays, it is required for effective RGI mitigation during plant–microbe interactions.

The transfer of the *iad* pathway into *Polaromonas* MF047 and *Paraburkholderia* MF376 demonstrates that IAA-degradative activity can be reconstituted in root-associated bacterial hosts beyond native *Variovorax*. In both chassis strains, engineered *iad* expression conferred IAA degradation and mitigated auxin-mediated RGI across multiple microbial contexts, with MF376 showing particularly robust activity, including in a *Pseudomonas* MF048-induced RGI context in which CL014 was ineffective (Figs. 4 and 5c). These results support MF376 as a promising chassis for deploying auxin-degrading functions in plant-associated environments. Beyond restoring the core degradative function, these engineered strains also reveal that the fate of pathway end products can vary across recipient hosts. In CL014, anthranilic acid produced from IAA degradation can feed into tryptophan biosynthesis, whereas this metabolic connection was not detected in engineered MF047 or MF376 (Extended Data Fig. 6h, i). This suggests that, although pathway transfer can reproduce the core auxin-removal function, downstream metabolites may not be processed in the same way in recipient hosts. Anthranilic acid also did not affect *Arabidopsis* growth in the absence of bacteria or broadly affect bacterial growth under the tested conditions (Extended Data Fig. 6j, k). Instead, its modest growth-promoting effect was observed only when *Arabidopsis* was treated with CL028 alone or with CL028 together with MF376::*marR73 iadC–K2*. These results suggest that anthranilic acid may participate in specific plant–microbe interactions rather than acting as a general plant- or bacterial-growth-promoting metabolite.

16S rRNA amplicon sequencing revealed no significant or consistent differences in broad root-community composition between chassis strains (CL014 ΔHS33, MF047, or MF376) and their corresponding *iad*-engineered derivatives under the tested conditions (Fig. 6b). In contrast, larger community-level differences were observed among the bacterial chassis, which differed markedly in their relative abundance, with MF376 reaching the highest abundance in both wild-type and engineered backgrounds (Fig. 6b, c; Extended Data Fig. 8 and Fig. 9). In a separate natural-soil assay, MF376 carrying the complete *iad* pathway significantly increased shoot and root biomass and root length relative to uninoculated controls and wild-type MF376 (Fig. 7 and Extended Data Fig. 10). These results highlight the importance of chassis selection and identify MF376 carrying the complete pathway as the most effective engineered strain under the conditions tested.

Together, this study advances our mechanistic understanding of microbial auxin metabolism and provides a platform for transferring mechanistically defined plant-beneficial functions into new root-associated bacterial chassis. In the future, this approach may inform the development of next-generation microbial bioinoculants to enhance plant performance in diverse soil environments.

## Methods

### Engineering bacterial strains

#### Knock-out mutant construction

Gene deletions in *V. paradoxus* CL014 were generated using the suicide vector pMo130, following previously established methods^9,12,37^. The vector backbone was PCR amplified and treated with DpnI to remove the template DNA. Primers used for mutant construction are listed in Supplementary Table 2. Upstream and downstream flanking regions of target genes were amplified from *V. paradoxus* CL014 genomic DNA using Platinum SuperFi II PCR Master Mix (Thermo Fisher Scientific). PCR products were purified using the DNA Clean & Concentrator Kit (Zymo Research) and assembled into pMo130 using HiFi Gibson Assembly Master Mix (New England Biolabs). Assembled plasmids were transformed into *E. coli* NEB 5-alpha (New England Biolabs), selected on Lysogeny Broth (LB, Thermo Fisher Scientific, BP1427) agar (2% w/v) supplemented with kanamycin (50 µg/mL), and verified by Sanger sequencing (Genewiz). Sequence-confirmed plasmids were transferred into the diaminopimelic acid (DAP) auxotrophic *E. coli* WM3064 for conjugation. Transformants were selected on LB agar containing kanamycin (50 µg/mL) and DAP (0.3 mM), and grown at 37 °C for 24 h. For biparental mating, donor *E. coli* WM3064 and recipient *V. paradoxus* CL014 (pre-grown in 50% TSB (Tryptic Soy Broth, Thermo Fisher Scientific, CM0129) agar (2% w/v) with ampicillin (100 µg/mL)) were washed twice with 50% TSB, mixed at a 1:1 volume ratio, pelleted (5,000 × *g*, 5 min), resuspended in 1/10 volume of 50% TSB, and spotted onto 50% TSB agar containing DAP (0.3 mM). After overnight incubation at 28 °C, exconjugants were selected on 50% TSB agar containing ampicillin and kanamycin without DAP, and incubated for 3–4 days. Resulting colonies were re-streaked onto fresh antibiotic 50% TSB agar to ensure clonality and remove residual donor cells. Single colonies were screened by colony PCR to confirm single-crossover integration. Verified integrants were cultured in 50% TSB with ampicillin and Isopropyl β-D-1-thiogalactopyranoside (IPTG, 1 mM). Cultures were plated on sucrose counter-selection agar (10 g/L tryptone, 5 g/L yeast extract, 10% w/v sucrose, 2% w/v agar, 100 µg/mL ampicillin, 1 mM IPTG) and incubated at 28 °C for 3-4 days. Resulting colonies were passaged in the same liquid medium, and deletion events were verified by PCR. Final deletion strains were re-streaked on 50% TSB with ampicillin and confirmed by diagnostic PCR using one primer located outside the deletion region and one within the deleted gene to ensure complete excision and strain purity. DNA templates for all PCRs were prepared using the following rapid lysis protocol^38^. A single colony or 6 µl of bacterial culture was mixed with 10 µl of alkaline lysis buffer (25 mM NaOH, 0.2 mM Na₂-EDTA, pH 12), incubated at 95 °C for 30 min, and neutralized with 10 µl Tris-HCl (40 mM, pH 7.5). The resulting material was used directly as PCR template (1:10, template:total PCR reaction volume).

#### Overexpression mutant construction

*V. paradoxus* CL014 *marR73 iadCDE* and *marR73 iadC-K2* were cloned into the broad-host-range vector pBBR1MCS-2 as previously described^9,12,39^, genomic fragments were amplified using Platinum SuperFi II PCR Master Mix and assembled into pBBR1MCS-2 via Gibson assembly using HiFi Gibson Assembly Master Mix (New England Biolabs). Primers used for mutant construction are listed in Supplementary Table 2. Circular template DNA was digested with DpnI, and assembled plasmids were transformed into *E. coli* NEB 10-beta (New England Biolabs). Transformants were selected on LB agar containing kanamycin (50 µg/mL), and plasmids were extracted (ZR Plasmid Miniprep Kit, Zymo Research) and verified by Sanger sequencing. Verified constructs were introduced into *V. paradoxus* CL014 ΔHS33 by tri-parental mating, using *E. coli* pRK2013 as a helper strain. Donor and helper strains were grown in LB with kanamycin (50 µg/mL) at 37 °C, and the recipient strain was cultured in 50% TSB with ampicillin (100 µg/mL) at 28 °C. All strains were pelleted (5,000 × *g*, 5 min), washed in 50% TSB twice, mixed at equal volumes, and spotted onto 50% TSB agar for overnight conjugation at 28 °C. Exconjugants were selected on 50% TSB agar supplemented with kanamycin (50 µg/mL) and ampicillin (100 µg/mL), confirming successful plasmid transfer into *V. paradoxus* CL014 ΔHS33.

#### Knock-in mutant construction

Two gene combinations, *marR73 iadCDE* and *marR73 iadC-K2*, were integrated into the *iad* locus of *V. paradoxus* CL014 ΔHS33, as well as three intergenic genomic sites in *Polaromonas* MF047 (IMG genome ID 2636416056 with insertion positions between Gene IDs: 2639079279–80, 2639079819–20, and 2639080354–55) and one site in *Paraburkholderia* MF376 (IMG genome ID 2521172625 with insertion position between Gene IDs 2521671121–22), using the pMo130 suicide vector (Extended Data Fig. 6b). Gene amplification, vector assembly, and verification followed the same procedure as knockout mutant construction, except that plasmids were initially propagated in *E. coli* NEB 10-beta. Primers used for mutant construction are listed in Supplementary Table 2. Sequence-verified plasmids were electroporated into *V. paradoxus* CL014 ΔHS33, *Polaromonas* MF047, and *Paraburkholderia* MF376, as described below. Strains were cultured in 50% TSB at 28 °C with shaking (250 rpm) for 2 days, followed by 24 h incubation at 4 °C. Cells were harvested (5,000 × *g*, 10 min, 4 °C), washed twice with ice-cold sterile water, and resuspended in 10% sterile glycerol for electroporation. For each reaction, 100 ng of plasmid DNA was electroporated into 100 µl of competent cells using a 0.1 cm gap cuvette using the following conditions: 1,800 V, 25 µF, 200 Ω for *marR73 iadCDE*, and 2,500 V, 25 µF, 200 Ω for *marR73 iadC-K2*. After electroporation, cells were recovered in SOC medium (New England Biolabs, B9020) at 28 °C (250 rpm) for 3 h and plated on 50% TSB agar with selective antibiotics: ampicillin (100 µg/mL) + kanamycin (200 µg/mL) for *Polaromonas* MF047 and kanamycin (50 µg/mL) for *Paraburkholderia* MF376. After 4–5 days of incubation, single colonies were screened by colony PCR using crude DNA extracted with alkaline lysis buffer. PCR-confirmed integrants were grown overnight in 50% TSB with 1 mM IPTG, supplemented with ampicillin (100 µg/mL) for *Polaromonas* MF047 and without antibiotics for *Paraburkholderia* MF376. Cultures were then diluted 1,000-fold and plated on sucrose counter-selection agar to induce second recombination (5% w/v sucrose for *Polaromonas* MF047; 20% w/v sucrose for *Paraburkholderia* MF376). Double-crossover mutants were confirmed by PCR and Sanger sequencing.

### Liquid chromatography-mass spectroscopy (LC–MS) metabolomics

#### Sample preparation

Strains were streaked from glycerol stocks onto 50% TSB agar plates supplemented with the appropriate antibiotics: ampicillin (100 µg/mL) for the CL014 wild type and knockout mutants, ampicillin (100 µg/mL) plus kanamycin (50 µg/mL) for CL014 overexpression strains, and no antibiotics for MF047- and MF376-derived strains. Plates were incubated at 28 °C for 3 days. Single colonies were inoculated into 5 mL of 50% TSB containing the appropriate antibiotics and cultured at 28 °C with shaking at 250 rpm for 48 h. Bacterial cultures were harvested and washed following the root growth inhibition assay protocol. CL014- and MF376-related strains were resuspended in 5 mL of modified M9 medium supplemented with 15 mM succinic acid, whereas MF047-related strains were resuspended in 50% TSB medium. In all cases, cultures were adjusted to a final OD₆₀₀ of 0.05. Cultures were then incubated at 28 °C with shaking at 250 rpm for 15 h for CL014- and MF376-related strains, or for 20 h for MF047-related strains, after which IAA or deuterium-labeled IAA ([²H₇]IAA, DLM-8040-0.1, Cambridge Isotope Laboratories, dissolved in 100% ethanol) was added to a final concentration of 0.1 mg/mL. Cultures were incubated for an additional 4 h for CL014- and MF376-related strains, or 12 h for MF047-related strains. For H₂¹⁸O labeling experiments, H₂¹⁸O (98%; ABX Advanced Biochemical Compounds GmbH, #894) was used to replace 20% of the water in the culture medium (M9). Cells were collected at a total biomass of OD₆₀₀ × volume (mL) = 2, centrifuged at 5,000 × *g* for 10 min at 4°C, and pellets were resuspended in 400 µl of ice-cold quenching solvent (acetonitrile : methanol : water, 40:40:20, v/v/v). Samples were stored at −80 °C prior to metabolite extraction and LC–MS/MS analysis.

#### LC-MS analysis

LC-MS analysis was performed on a Vanquish UHPLC system (Thermo Fisher Scientific) coupled to a quadrupole Orbitrap Exploris 480 mass spectrometer. LC separation of polar metabolites was achieved using a Waters XBridge BEH Amide column (2.1 mm × 150 mm, 2.5-µm particle size, 130-Å pore size). The LC method used a 25-min solvent gradient at a flow rate of 150 µL/min, with the following gradient parameters: 0 min, 90% B; 2 min, 90% B; 3 min, 75% B; 7 min, 75% B; 8 min, 70% B, 9 min, 70% B; 10 min, 50% B; 12 min, 50% B; 13 min, 25% B; 14 min, 25% B; 16 min, 0% B, 20.5 min, 0% B; 21 min, 90% B; 25 min, 90% B, where Solvent A was 95:5 water : acetonitrile with 20 mM ammonium hydroxide and 20 mM ammonium acetate (pH 9.4) and solvent B was acetonitrile. The autosampler temperature was 4 °C, the column temperature was 25 °C, and the injection volume was 10 μl. N-formylanthranilic acid standard was obtained from AldrichCPR (S636886). The Exploris 480 mass spectrometer was operated in full scan mode in negative polarity on MS1 level, which allows the relative quantitation of the metabolite across by ion count. Following parameters are used for the full scan: resolution, 120,000; scan range, m/z 70-1000 (negative mode); AGC target, 1e6; ITmax, 500 ms. Other instrument parameters are spray voltage 3000 V, sheath gas 35 (Arb), aux gas 10 (Arb), sweep gas 0.5 (Arb), ion transfer tube temperature 300 °C, vaporizer temperature 35◦C, internal mass calibration on, RF lens 50. The MS2 spectra were collected in targeted mode using the parallel reaction monitoring (PRM) function at higher energy C-trap dissociation (HCD) energy of 20 eV, and other instrument settings as following: resolution 30,000, AGC target 1e6, maximum injection time 250 ms, and isolation window 1.0 m/z. For the MS1 data analysis, raw LC–MS data were converted to mzXML format using *ProteoWizard*^40^ Peak picking was performed with EL-Maven (v0.12.1-beta; Elucidata) for unlabeled and ^2^H-labeled compounds, and MAVEN (v2.10.14c) for ^18^O-labeled compounds. Relative abundance changes of each metabolite were quantified using relative peak area tops in the chromatogram. For ^2^H-labeled data analysis, natural isotope abundance was corrected using the AccuCor R package^41^ (https://github.com/lparsons/accucor). For ^18^O-labeled data analysis, isotope correction was performed using the Iso-Autocorr package (https://github.com/xxing9703/Iso-Autocorr). For the MS2 data, raw LC–MS files were processed and peaks were extracted with the built-in Xcalibur Qual Browser (Thermo Scientific, v4.4).

#### Synthesis and characterization of C₁₀H₉NO₃

The target compound was synthesized following a reported procedure with modifications^42^. Briefly, indole-3-acetic acid (48 mg) was dissolved in acetone (11 mL), followed by addition of saturated aqueous sodium bicarbonate (14 mL) at room temperature. An aqueous solution of Oxone (213 mg in 4 mL water) was added dropwise over 10 min. The reaction mixture was stirred for an additional 10 min and then filtered. Acetone was removed under reduced pressure using a rotary evaporator, and the remaining aqueous phase was lyophilized. The residue was redissolved in methanol (5 mL), centrifuged at 12,000 × *g* for 5 min, and the supernatant was subjected to preparative HPLC purification. Preparative HPLC was performed on an Agilent 1260 system equipped with an Agilent ZORBAX StableBond C18 column (9.4 × 250 mm, 5 µm). Buffer A consisted of water containing 0.1% formic acid, and buffer B consisted of acetonitrile containing 0.1% formic acid. The purification was carried out with a 1 mL injection volume and a flow rate of 10 mL min⁻¹ using the following gradient: 0–2.5 min, 90% A; 2.5–7.5 min, linear gradient from 90% A to 60% A; 7.5–12.5 min, linear gradient from 60% A to 5% A; 12.5–15 min, 5% A. Fractions were analyzed by LC–MS, and pure fractions were pooled and concentrated by lyophilization. The purified product was obtained as a white powder and redissolved in CD₃CN for NMR and IR analysis. Based on the proposed epoxide ring-opening mechanism, the synthesized lactone is expected to adopt a *cis*-fused configuration at the lactone junction.

NMR spectra were collected on a Bruker Advance II spectrometer operating at 500 MHz and at 298 K, equipped with a cryoprobe at Princeton University. ^1^H NMR (500 MHz, CD₃CN) δ 7.32 (dd, *J* = 7.4, 1.3 Hz, 1H), 7.20 (td, *J* = 7.7, 1.3 Hz, 1H), 6.85 (td, *J* = 7.5, 1.0 Hz, 1H), 6.73 (d, *J* = 7.9 Hz, 1H), 5.86 (br, 1H), 5.73 (s, 1H), 3.11 (d, *J* = 17.8 Hz, 1H), 3.08 (d, *J* = 17.8 Hz, 1H); ^13^C NMR (126 MHz, CD₃CN) δ 174.45, 149.07, 131.61, 130.88, 125.41, 120.81, 111.07, 100.19, 83.38, 42.12. IR spectra were collected on Nicolet iS50 FTIR Spectrometer (Thermo Fisher Scientific). IR (ATR, neat, cm^-1^): 3380, 1763, 1614, 1474, 1315, 1204, 1104, 934, 749. The NMR and IR spectra are provided in the Supplementary Information.

### Cell lysate reaction

A 50 mL culture of ΔHS33_p_*iadCDEJK2* was prepared and harvested by centrifugation at 5,000 × *g* for 10 min at 4℃. The resulting cell pellet was resuspended in 1 mL of reaction buffer (100 mM HEPES, 50 mM NaCl, pH 7.5) and lysed by sonication using a Qsonica Sonicator 3000 (total sonication time, 1 min; 50% amplitude; 5 s on, 25 s off). After centrifugation at 20,000 × *g* for 10 min, the supernatant was collected and used directly for cell lysate reactions without further purification. Each 100 µL reaction contained 50 µL of cell lysate, 1 mM synthesized substrate (C₁₀H₉NO₃), and 2 mM NAD⁺ or NADP⁺ in reaction buffer. To quench the reaction, a 5 µL aliquot was mixed with 35 µL methanol, followed by centrifugation at 20,000 × *g* for 10 min. The resulting supernatant was then subjected to LC–MS analysis.

### IadDE protein expression, purification and crystallization

#### Protein expression

The *iadD* and *iadE* genes (IMG gene IDs: 2643613669, 2643613668) from *V. paradoxus* CL014 (IMG genome ID: 2643221508) were cloned into the pET28b expression vector with an N-terminal His-tag and transformed into *E. coli* Rosetta(DE3) (Millipore Sigma) for recombinant protein expression. Transformants were plated on LB agar supplemented with kanamycin (50 µg/mL) and chloramphenicol (33 µg/mL) and incubated at 37 °C for 24 h. Individual colonies were picked and grown overnight in LB medium at 37 °C with shaking at 250 rpm. Overnight cultures were diluted 1:1000 into autoinduction (AI) medium (ZYM-5052)^43^ containing the same antibiotics, and incubated at 37 °C, 250 rpm for 22 h. Cells were harvested by centrifugation at 5,000 × *g* for 20 min at 4 °C, and pellets were collected and stored at –80 °C for subsequent protein purification.

#### Protein purification

Cell pellets were resuspended in IMAC Buffer A (20 mM NaH₂PO₄, 500 mM NaCl, pH 7.4) at a ratio of 1 g pellet per 10 mL buffer. To reduce viscosity, 0.1 µL benzonase nuclease (Sigma-Aldrich, 70746) was added per gram of cell pellet. Cells were lysed using an EmulsiFlex-C5 high-pressure homogenizer (three passes), and the lysate was clarified by centrifugation at 25,000 × *g* for 30 min at 4 °C. The supernatant was filtered through a 0.22 µm syringe filter and loaded onto a 5 mL Ni-charged Nuvia IMAC column (Bio-Rad) using a fast protein liquid chromatography (FPLC) system (Bio-Rad). The column was washed with IMAC Buffer A containing 15 mM imidazole, and bound proteins were eluted with IMAC Buffer B (20 mM NaH₂PO₄, 500 mM NaCl, 500 mM imidazole, pH 7.4). Eluted fractions were analyzed by SDS–PAGE (Bio-Rad Stain-Free gels), pooled, concentrated, and buffer-exchanged into 50 mM Tris-HCl (pH 7.0), 150 mM NaCl using 10 kDa MWCO concentrators (Pierce, Thermo Fisher). Purified protein was stored at 4 °C, and concentration was determined using the Pierce BCA Protein Assay Kit (Thermo Fisher Scientific).

#### Protein crystallography

Purified IadDE was concentrated to 10 mg/mL in buffer supplemented with 1 mM dithiothreitol (DTT) and crystallized using the sitting-drop vapor diffusion method at 20 °C. Crystallization drops were prepared by mixing 1.5 µl of protein solution with 1.5 µl of reservoir solution in 24-well sitting-drop plates. The reservoir solution contained 9.6–10.6% (w/v) PEG 3350 and 0.1 M sodium citrate tribasic dihydrate (pH 5.5). Crystals were cryoprotected by brief soaking in the reservoir solution supplemented with 30% (v/v) ethylene glycol and subsequently flash-cooled in liquid nitrogen for data collection. Diffraction data were collected at beamlines 17-ID1 (AMX) and 17-ID2 (FMX) at Brookhaven National Laboratory to a maximum resolution of 1.28 Å. Crystals grew in space group H3 (hexagonal setting of R3) with typical cell dimensions a=b=130.5 Å c=100.7 Å α=β=90^°^ γ=120^°^ with one complex per asymmetric unit. Data were processed with XDS^44^ and scaled with AIMLESS^45^. The structure was determined by the method of molecular replacement using the program PHASER^46^ utilizing sequential placement of the two subunits with the models derived from AlphaFold models. The structure was rebuilt in COOT^47^, incorporating a [2Fe–2S] cluster and a mononuclear iron-binding site, and subsequently refined using PHENIX.REFINE^48^. The final model showed good agreement with the experimental data and displayed excellent geometry. The final model and associated X-ray data have been deposited with the Protein Data Bank with code 9O71. Relevant refinement statistics are summarized in Supplementary Table 3.

### Measurement of IAA degradation

IAA degradation by bacterial strains was assessed using a spike-in approach. Bacterial preparation followed the same procedure as the root growth inhibition assay, including streaking from glycerol stocks, cultivation, washing, and OD₆₀₀ measurement. Washed cells were inoculated to a final OD₆₀₀ of 0.05 into either M9 medium^9,12^(3 g L⁻¹ KH₂PO₄, 0.5 g L⁻¹ NaCl, 6.78 g L⁻¹ Na₂HPO₄, and 1 g L⁻¹ NH₄Cl) supplemented with 2 mM MgSO₄, 0.1 mM CaCl₂, 10 µM FeSO₄, and 5 g L⁻¹ glucose for CL014- and MF376-related strains, or 50% TSB medium for MF047-related strains. Cultures were incubated at 28 °C with shaking (250 rpm) for 15 h before IAA (dissolved in 100% ethanol) was spiked-in to a final concentration of 0.1 mg/mL. Aliquots (300 µl) were collected at 2 h or 4 h intervals, centrifuged at 5,000 × *g* for 10 min, and 50 µl of the supernatant was mixed with 100 µl of freshly prepared Salkowski reagent (10 mM FeCl₃ and 35% perchloric acid). After incubation for 40 min at room temperature, absorbance was measured at 530 nm using a BioTek Synergy H1 microplate reader. As *Polaromonas* MF047 is an amino acid dependent strain and cannot grow in M9 medium, this strain and its engineered strains were assayed in 50% TSB medium instead of M9. All strains were tested in three biological replicates to ensure reproducibility.

### Bacterial growth curve

Bacterial preparation followed the same protocol as in the root growth inhibition assay. Washed cells were inoculated into 200 µl of 50% TSB medium supplemented with or without IAA to a final OD₆₀₀ of 0.05. Each strain was tested in three biological replicates to ensure reproducibility. Cultures were grown in sterile 96-well cell culture plates, sealed with Breathe-Easy gas-permeable film (Diversified Biotech), and incubated at 28 °C with continuous linear shaking. Optical density at 600 nm (OD₆₀₀) was recorded every 2 h over a 24-hour period using a BioTek Synergy H1 microplate reader to monitor bacterial growth dynamics.

### Root growth inhibition assay

#### Seedling and seed preparation

*Arabidopsis thaliana* Col-0 seeds were surface-sterilized by vortexing in 70% bleach containing 0.2% Tween-20 for 10 min, followed by five washes with sterile distilled water. Seeds were sown on half-strength Murashige and Skoog (MS) agar medium (2.22 g/L MS basal medium with Gamborg vitamins (PhytoTech Labs, M-404), 0.5 g/L MES, 5 g/L sucrose, 10 g/L agar, pH 5.7 adjusted with 3 M NaOH) in 12 × 12 cm square plates and grown vertically under short-day conditions (21 °C day / 18 °C night, 10 h light / 14 h dark, 70% relative humidity, 170 μmol m⁻² s⁻¹ light intensity) for 7 days. *Medicago* seeds were sterilized following published protocols^49,50^. Briefly, seeds were treated with concentrated sulfuric acid for 10 min with agitation, rinsed once with sterile water, then treated with 70% bleach for 3 min. After five additional washes, seeds were imbibed in sterile water for 2–6 h at room temperature prior to use.

#### IAA or anthranilic acid-containing plate preparation

Half-strength MS agar medium was autoclaved and cooled until warm to the touch. IAA stock solution (dissolved in 100% ethanol) or anthranilic acid stock solution (dissolved in water) was added to the final concentration noted. Stock solutions were stored at –20 °C for up to two months.

#### Bacterial and SynCom32 preparation

Bacterial strains were revived from 20% glycerol stocks by streaking onto 50% TSB agar plates (15 g/L tryptic soy broth, 20 g/L agar) and incubated at 28 °C for 3–4 days. Single colonies were inoculated into 5 mL 50% TSB and grown for 2 days at 28 °C with shaking (250 rpm). Cultures were pelleted at 5,000 × g for 10 min, washed twice with 3 mL of sterile 10 mM MgCl₂, and resuspended in 750 µl of the same buffer. The optical density at 600 nm (OD₆₀₀) was measured using a NanoDrop One C with semi-micro cuvettes and adjusted to 0.05. For SynCom32 assembly, each strain’s OD₆₀₀ was measured individually. To ensure equal biomass contribution, volumes were calculated using the equation: OD₆₀₀ × volume (µl) = 300, and combined accordingly. The pooled culture was washed and resuspended as described above.

#### Bacterial inoculation

For plant-microbe co-inoculation assays, 100 µl of strain (OD₆₀₀ = 0.05) was evenly spread onto the surface of half-strength MS agar plates. For SynCom32, 100 µl of the pooled culture (OD₆₀₀ = 0.05) was applied, followed by 100 µl of *V. paradoxus* CL014, *Polaromonas* MF047, *Paraburkholderia* MF376, or their engineered strains at OD₆₀₀ = 0.005. Plates were incubated overnight at room temperature prior to seedling transfer.

#### Plant growth and measurement

Seven-day-old *Arabidopsis* seedlings or sterilized *Medicago* seeds were transferred onto pre-inoculated plates. Plates were sealed with 3M Micropore tape and incubated vertically in a growth chamber under short-day conditions for 7 days. For 16S rRNA amplicon sequencing, plants were grown for 9 days to allow sufficient biomass for DNA extraction. Plates were imaged using a digital camera, and primary root elongation was measured as the distance from the initial to final root tip position using the freehand line tool in ImageJ. All primary root elongation data are provided in the Source Data file.

### Microbiome analysis

#### Sample collection

After nine days of co-incubation with bacteria, roots from *Arabidopsis* and *Medicago* were harvested for microbiome profiling. Each sample consisted of 5–10 roots pooled per plate, with 3–5 biological replicates per treatment. Roots were transferred to 15 mL Falcon tubes containing 7 mL sterile water and washed by vigorous vortexing. Excess water was removed using sterile filter paper, and dried roots were transferred to Lysing Matrix E tubes (MP Biomedicals) and stored at –80 °C until DNA extraction. For SynCom32 input controls, bacterial cultures were pelleted at 5,000 × *g* for 10 min, the supernatant was discarded, and pellets were stored at –80 °C.

#### DNA extraction

Samples were homogenized using a FastPrep-24™ 5G instrument (MP Biomedicals) with two 40 s cycles at 6.0 m/s for root samples, and one cycle for SynCom32 input controls, with 2 min on ice between runs to prevent overheating. DNA was extracted using the FastDNA SPIN Kit for Soil (MP Biomedicals), eluted in 55 µl DES buffer (provided in the kit), quantified using the Quant-iT PicoGreen dsDNA Assay Kit (Thermo Fisher), and diluted to 3.5 ng/µl with DES buffer.

### 16S rRNA library preparation and sequencing

The V5–V7 region^51^ of the 16S rRNA gene was amplified using a two-step dual-indexed PCR approach. The first PCR was performed using Platinum SuperFi II PCR Master Mix (Thermo Fisher) in a 25 µl reaction containing: 12.5 µl Master Mix, 7 µl nuclease-free water (Qiagen), 2 µl of 5 µM forward primer 799F (5′-AACMGGATTAGATACCCKG-3′, with 10-bp sample barcode), 1 µl of 10 µM reverse primer 1192R (5′-ACGTCATCCCCACCTTCC-3′, with 6-bp library barcode), and 2.5 µl of 3.5 ng/µl DNA template. Thermal cycling conditions were: 98 °C for 30 s, followed by 30 cycles of 98 °C for 10 s, 55 °C for 10 s, and 72 °C for 15 s, with a final extension at 72 °C for 5 min. Three technical PCR replicates were pooled per sample to minimize amplification bias. Products were verified by 1.2% agarose gel electrophoresis. For gel purification, 25 µl of each PCR product (two samples per lane) were pooled with 10 µl 6× loading dye, run on a 1.2% agarose gel, and ∼400 bp bands were excised and purified using the Wizard SV Gel and PCR Clean-Up System (Promega). DNA concentration was assessed by PicoGreen, and 100 ng of each sample was pooled for library construction and cleaned with 0.9× AMPure XP beads (Beckman Coulter). A second PCR was performed to add Illumina sequencing adapters. Final libraries were sequenced on an Illumina MiSeq platform (2 × 300 bp paired-end reads)^51,52^ at the Princeton Genomics Core Facility. After quality filtering, a total of 14,907,735 high-quality sequences were obtained from 175 samples, with an average of 85,187 reads per sample.

#### Amplicon data processing

Raw reads were demultiplexed in QIIME2 (v2024.10)^53^ using qiime cutadapt demux-paired^54^, and primer sequences were trimmed with qiime cutadapt trim-paired. Denoising, quality filtering, and chimera removal were performed with DADA2 (qiime dada2 denoise-paired)^55^. After testing multiple truncation lengths, forward reads were truncated at 220 bp and reverse reads at 180 bp to optimize read retention and ASV diversity. ASVs were taxonomically classified using a naïve bayes classifier trained on a custom database of root-associated bacterial sequences^56^ via qiime feature-classifier classify-sklearn. Assignments were filtered based on confidence scores, prevalence, and relative abundance. Relative abundance tables were generated using qiime feature-table relative-frequency and merged with taxonomy and metadata for visualization in R using ggplot2. Beta diversity was calculated using Bray–Curtis dissimilarity (R package vegan, vegdist)^57^, followed by principal coordinate analysis (PCoA; cmdscale). Differences in community composition were assessed by PERMANOVA using adonis2. Differential abundance analysis was conducted using the Mann–Whitney U test with FDR correction. Log₂ fold changes were calculated, and heatmaps were visualized using pheatmap (R v4.4.2)^58^.

### Plant growth promotion assay in natural soil

Soil was collected from the Stony Ford Research Station (Princeton, NJ, USA), where no chemical fertilizers, pesticides, or plants had been applied or grown in recent years. The soil was sieved twice to remove rocks and plant debris, then distributed into pots placed within 9 × 13-inch aluminum foil trays. Pots were saturated overnight with 1.2 L of sterile water containing bacterial inoculum. Bacterial preparation followed the same protocol as the root growth inhibition assay, including streaking, cultivation, washing, and OD₆₀₀ measurement. Individual bacterial strains were suspended in 1.2 L of sterile water to a final OD₆₀₀ of 0.03 prior to soil application. Sterile *Arabidopsis* and *Medicago* seeds were prepared using the same sterilization procedures described for the root growth inhibition assay. Pots were maintained in a growth chamber under short-day conditions and watered with 600–800 mL of distilled water every 4 days. Two sets of soil experiments were conducted. In the first set, each pot contained three to four *Arabidopsis* plants, and the experiment was designed specifically to measure shoot fresh weight. In the second set, smaller pots were used, with one plant per pot, to enable collection of multiple traits from the same individual plant, including shoot fresh weight, shoot dry weight, root fresh weight, root dry weight, and root length. This second set included both *Arabidopsis* and *Medicago*. After 33 days, plants were harvested. For *Arabidopsis*, roots grown in natural soil were too fragile to be reliably recovered intact, and therefore only shoot traits were obtained. For *Medicago*, the full set of shoot and root measurements was successfully collected. Roots were washed gently with water to remove adhering soil. Fresh weights were measured immediately after harvest. Samples were then wrapped in aluminum foil, dried at 65 °C for 7 days, and weighed again to determine dry weight.

## Acknowledgements

We thank Prof. Jeffery L. Dangl (University of North Carolina at Chapel Hill, USA) for providing bacterial strains used in this study and for valuable suggestions on the manuscript. We also acknowledge the Princeton University Genomics Core Facility for performing 16S rRNA amplicon sequencing. This research used resources of the AMX (17-ID-1) and FMX (17-ID-2) beamlines at the National Synchrotron Light Source II (NSLS-II), a U.S. Department of Energy (DOE) Office of Science User Facility operated by Brookhaven National Laboratory under Contract No. DE-SC0012704. The Center for Bio Molecular Structure (CBMS) is supported by the NIH National Institute of General Medical Sciences (P30GM133893) and the DOE Office of Biological and Environmental Research (KP1605010). This work was supported by the Lidow Independent Work/Senior Thesis Fund to M.J.K.; the National Science Foundation grant CHE-2246289 to J.T.G.; the Department of Energy (DOE) DE-SC0018260 to J.D.R; the DOE Center for Advanced Bioenergy and Bioproducts Innovation (U.S. Department of Energy, Office of Science, Biological and Environmental Research Program under Award Number DE-SC0018420) to J.D.R., Y.S., and X.L.; the Project X Fund administered by the School of Engineering and Applied Science at Princeton University to J.M.C.; startup funds from the Department of Chemical and Biological Engineering to J.M.C.

## Author contributions

J.M.C. and J.D.R. supervised the project. T.J., Y.S., and J.M.C. conceived the study and designed the experiments. T.J. constructed mutant bacterial strains and prepared samples for LC-MS analysis. Y.S. and X.L. conducted metabolomics experiments, and together with J.T.G., analyzed the resulting data. Y.Z. synthesized and purified the compound, performed the cell lysate experiments, and organized the associated data. T.J. performed plant-microbe interaction experiments and data analysis. T.J. and M.J.K. expressed and purified proteins, while T.J. and P.D.J. carried out protein crystallization and structural data analysis. T.J. also prepared samples, constructed libraries, and analyzed 16S rRNA sequencing data, and conducted the plant growth promotion assay in natural soil. T.J., Y.S., X.L., and J.M.C. wrote the manuscript, with input and feedback from all co-authors.

## Data availability

All data supporting the findings of this study are available within the paper and its Supplementary Information. Source data underlying the figures are also provided with this paper. The raw 16S rRNA gene amplicon sequencing reads generated in this study have been deposited in the NCBI Sequence Read Archive (SRA) under BioProject accession number PRJNA1466608. The associated BioSample and SRA accession numbers are available through the same BioProject. The IadDE protein structure has been deposited in the Protein Data Bank under accession code 9O71.

