## Extended Data Figures for "Engineering auxin degradation into root-associated bacteria promotes plant growth"

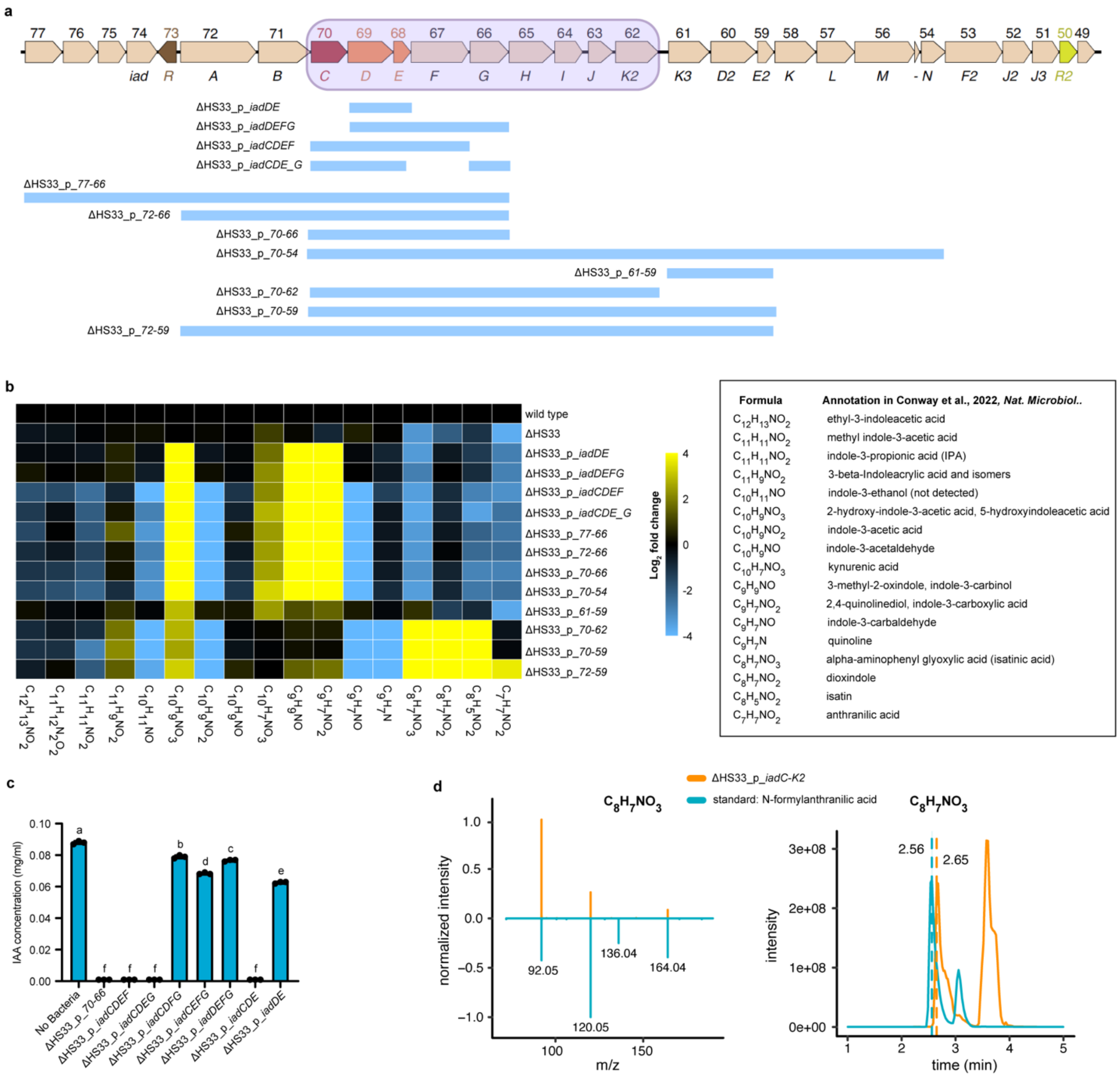


**Extended Data Fig. 1 | The *iadC-K2* gene region mediates IAA degradation in *V. paradoxus* CL014. a**, Cloned fragments from the *iad* locus genomic region inserted into the broad-host-range vector pBBR1 for functional validation in Hot Spot 33 (HS33) deletion strain *V. paradoxus* ΔHS33 from[9]. **b**, Heatmap showing $\log_2$ fold changes in metabolite abundance in complemented mutant strains relative to the wild type. Mass features (molecular formula and retention time in minutes) were selected based on previously reported IAA degradation intermediates[9]. **c**, Quantification of IAA concentration after 4 h of incubation in M9 minimal medium supplemented with 0.1 mg $ml^{-1}$ IAA. Data represent the mean ± s.d. of $n$ = 3 biological replicates. Statistical significance was assessed by one-way ANOVA followed by Tukey's post hoc test; different letters denote statistically distinct groups. **d**, LC–MS/MS validation showing that $C_8H_7NO_3$ is not N-formylanthranilic acid. Left,

MS/MS comparison of intracellular $C_8H_7NO_3$ and an authentic N-formylanthranilic acid standard, with fragment intensities normalized to the maximum signal within each sample. Right, extracted ion chromatograms showing distinct retention times.

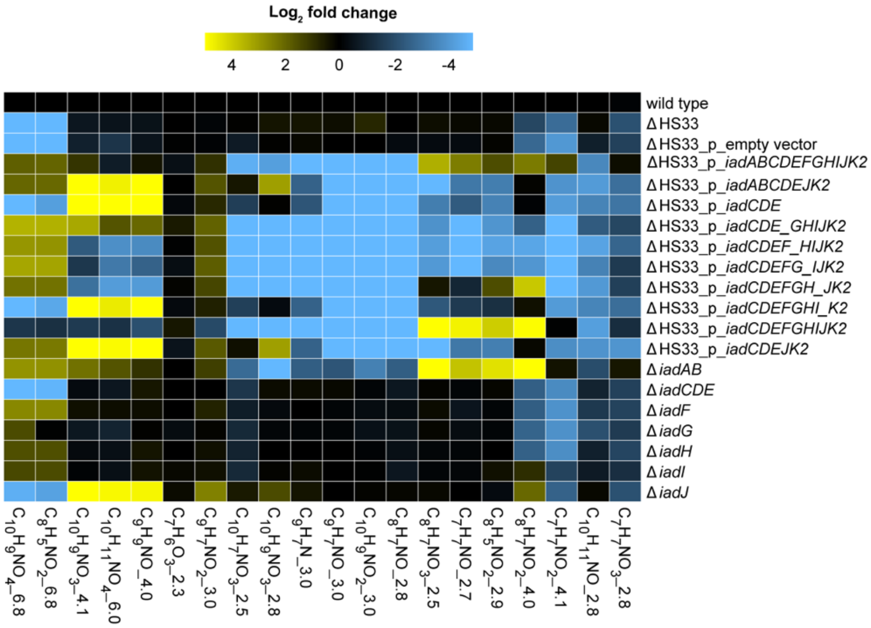


**Extended Data Fig. 2 | LC–MS analysis of *V. paradoxus* CL014 *iad* pathway mutants.** Each detected mass feature is annotated with its molecular formula and retention time (min). The heatmap displays $\log_2$ fold changes in metabolite abundance relative to wild-type CL014. This dataset supports metabolite assignments and pathway inference presented in Fig. 1b. Data represent the mean of $n$ = 3 biological replicates.

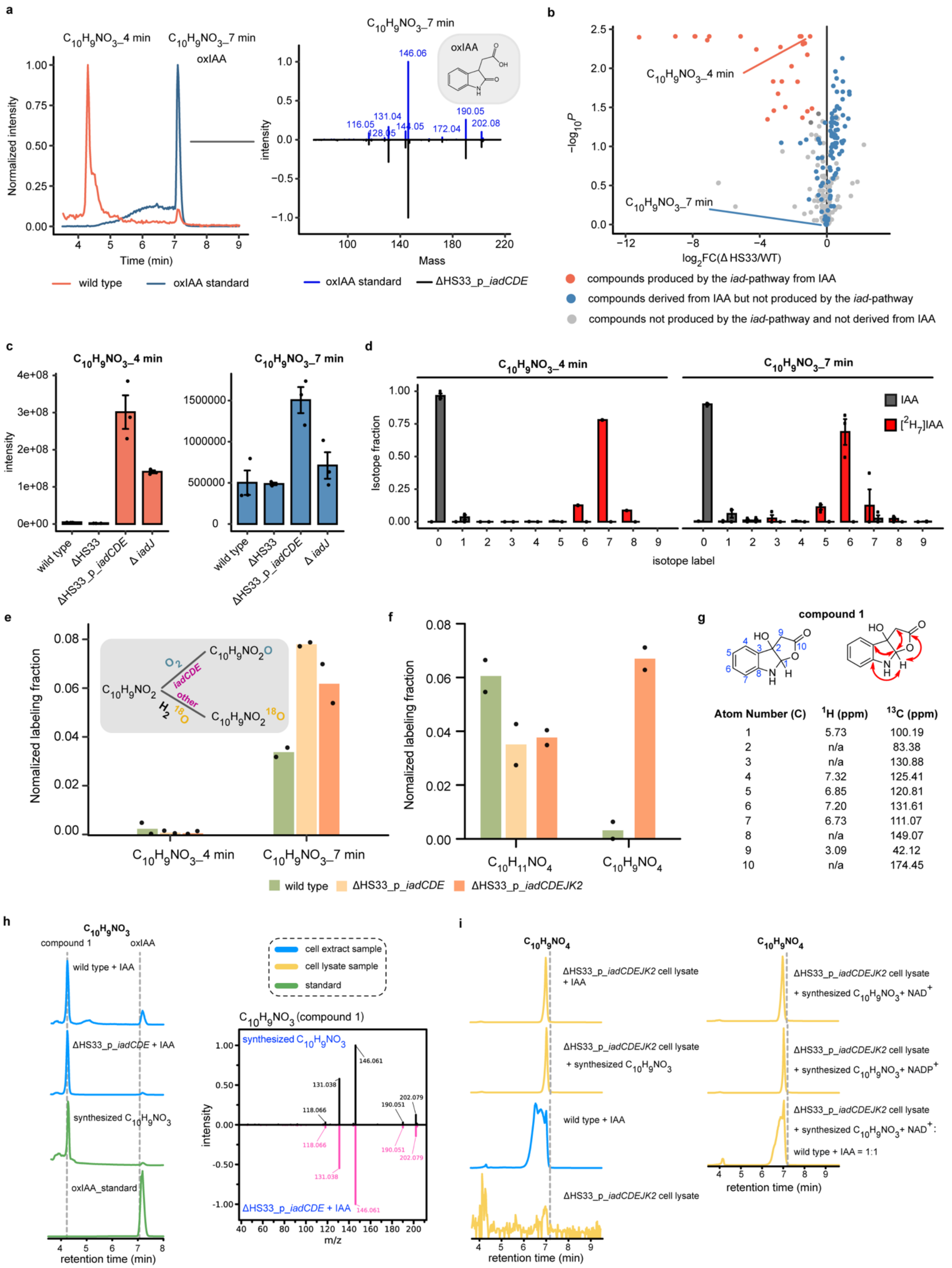


| Atom Number (C) | $^1$H (ppm) | $^{13}$C (ppm) |
|---|---|---|
| 1 | 5.73 | 100.19 |
| 2 | n/a | 83.38 |
| 3 | n/a | 130.88 |
| 4 | 7.32 | 125.41 |
| 5 | 6.85 | 120.81 |
| 6 | 7.20 | 131.61 |
| 7 | 6.73 | 111.07 |
| 8 | n/a | 149.07 |
| 9 | 3.09 | 42.12 |
| 10 | n/a | 174.45 |

**Extended Data Fig. 3 | Discovery of a $C_{10}H_9NO_3$ compound distinct from oxIAA during IadCDE-mediated IAA degradation. a**, LC–MS/MS analysis identified two distinct $C_{10}H_9NO_3$ compounds; comparison with a standard confirmed the 7 min peak as oxIAA. **b**, $Log_2$ fold changes in metabolite abundance in ΔHS33 relative to wild type. Statistical significance was assessed for *n* = 3 biological replicates using ANOVA followed by Benjamini–Hochberg correction for multiple comparisons. **c**, Relative abundance of the two $C_{10}H_9NO_3$ compounds in wild-type and mutant strains. Data represent the mean of *n* = 3 biological replicates. **d**, Deuterium isotope distribution profiles of $C_{10}H_9NO_3$ compounds following incubation with unlabeled IAA or $[^2H_7]$IAA. Values represent mean ± s.e.m. of *n* = 3 biological replicates. **e**,**f**, $^{18}O$ incorporation from $H_2{}^{18}O$ (20%, v/v) into IAA-derived intermediates measured by LC–MS. **e**, Normalized $^{18}O$ labeling fractions of the two $C_{10}H_9NO_3$ peaks. **f**, Normalized $^{18}O$ labeling fractions of $C_{10}H_{11}NO_4$ and $C_{10}H_9NO_4$. Data represent *n* = 2 biological replicates. **g–i**, Structural validation and downstream conversion of the rearranged $C_{10}H_9NO_3$ intermediate. **g**, NMR-based structural assignment of the synthesized $C_{10}H_9NO_3$ compound. Key supportive $^1H$–$^{13}C$ HMBC correlations are indicated by red arrows. Raw NMR spectrums are provided in the Supplementary Information. **h**, Extracted ion chromatograms and MS/MS spectra comparing synthesized $C_{10}H_9NO_3$ with the metabolite detected in wild type + IAA and ΔHS33_p_*iadCDE* + IAA. **i**, Cell lysate assays showing conversion of synthesized $C_{10}H_9NO_3$ to $C_{10}H_9NO_4$ by ΔHS33_p_*iadCDEJK2*. The y-axis indicates intensity normalized to the maximum signal within each sample.

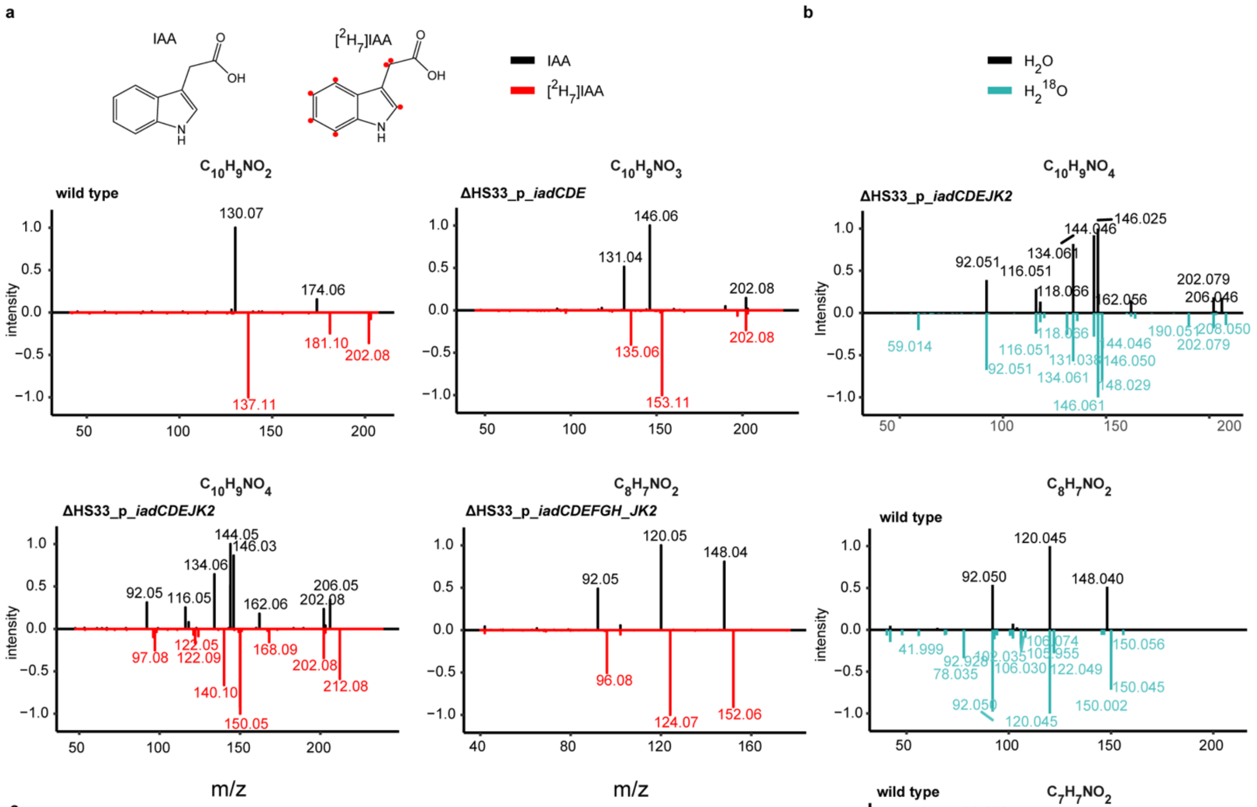


c

| unlabeled | # $^2$H | # $^{18}$O | | mass from IAA | mass from [$^2$H$_7$]IAA | transformation |
|---|---|---|---|---|---|---|
| C10H9NO2 | 7 | 0 | parent | 174.0562 | 181.0996 | |
| | 7 | 0 | | 130.0666 | 137.11 | -CO2 |
| C10H9NO3 | 7 | 0 | parent | 190.0508 | 197.0961 | |
| | 6 | 0 | | 160.0404 | 166.0779 | -CH2O |
| | 7 | 0 | | 146.061 | 153.1051 | -CO2 |
| | 4 | 0 | | 131.0376 | 135.0628 | -C2H3O2 |
| | 6 | 0 | | 118.0659 | 124.1042 | -CO -CO2 |
| | 5 | 0 | | 92.0504 | 97.0818 | C6H6N |
| C10H9NO4 | 6 | 1 | parent | 206.046 | 212.0835 | |
| | 6 | 0 or 1 | | 162.0561 | 168.0938 | -CO2 |
| | 4 | 0 or 1 | | 146.0251 | 150.0491 | -C2H4O2 |
| | 6 | 0 or 1 | | 144.0454 | 150.0831 | -CO2 -H2O |
| | 6 | 0 | | 134.0611 | 140.0988 | -CO2 -CO |
| | 6 | 0 | | 116.0505 | 122.0882 | -CO2 -CO -H2O |
| | 5 | 0 | | 92.0504 | 97.0819 | C6H6N |
| C8H7NO2 | 4 | 1 | parent | 148.0405 | 152.0656 | |
| | 4 | 0 or 1 | | 120.0454 | 124.0706 | -CO |
| | 4 | 0 | | 92.0506 | 96.0756 | C6H6N |

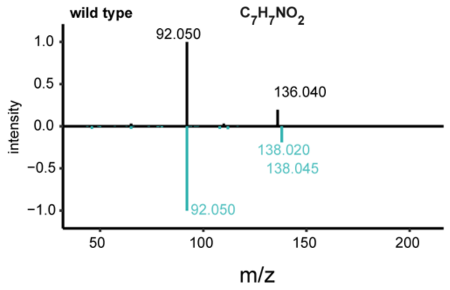


**Extended Data Fig. 4 | Isotope tracing and LC-MS/MS analyses reveal atom-level transformations in the IAA degradation pathway intermediates. a**, MS/MS spectra of major pathway intermediates following incubation with IAA or [$^2H_7$]IAA. **b**, MS/MS spectra of major pathway intermediates with $H_2O$ or $H_2{}^{18}O$ (20%, v/v). **c**, Summary table of parent ion masses, retained deuterium and $^{18}O$ atoms, and major mass fragment transformations across detected intermediates.

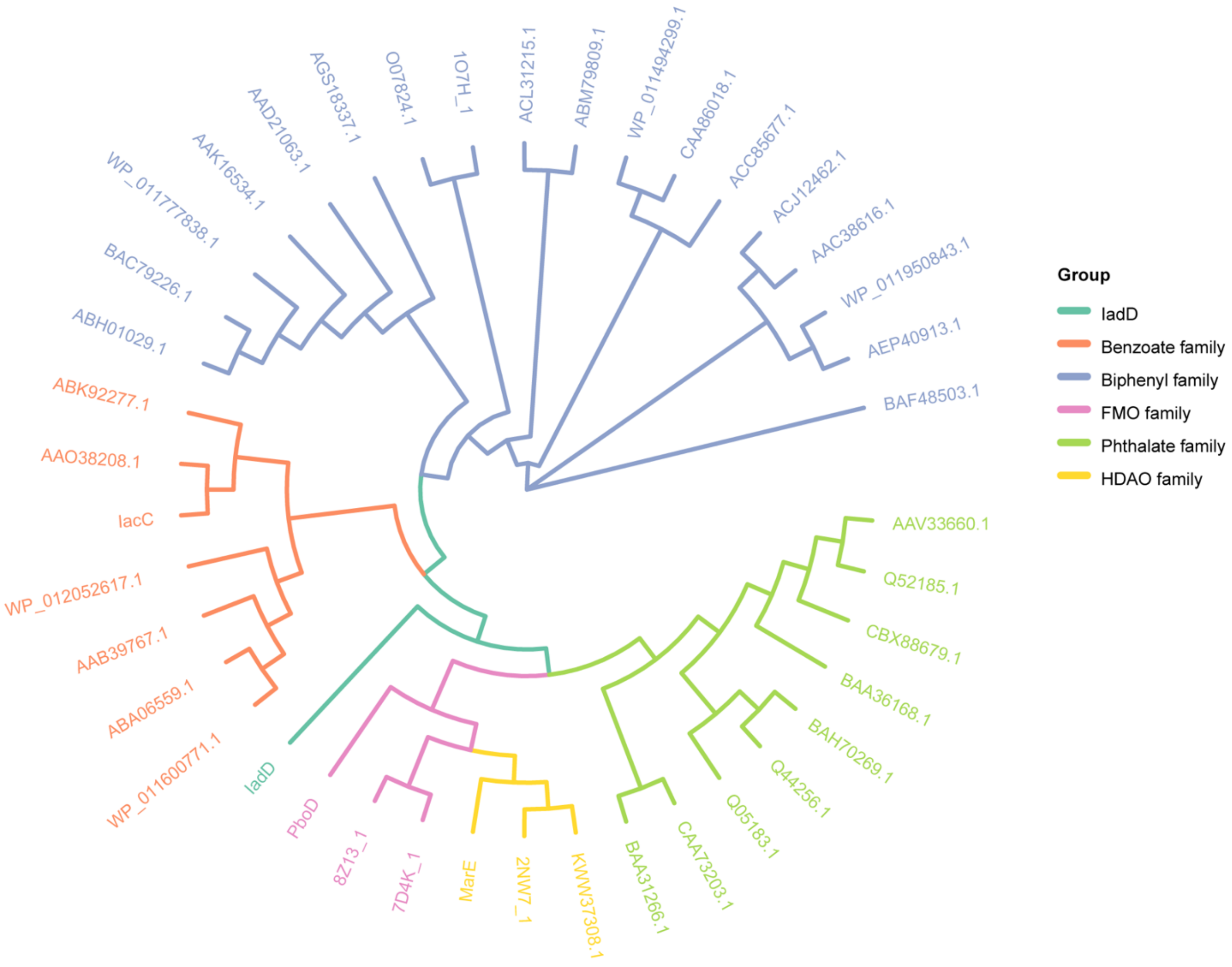


**Extended Data Fig. 5 | Phylogenetic analysis of IadD with Rieske dioxygenases and other indole oxygenases.** Protein sequences of characterized Rieske non-heme dioxygenases, heme-dependent aromatic oxygenases (HDAOs), and flavin-dependent monooxygenases (FMOs) were obtained from NCBI and PDB (see Supplementary Table 5). Clustal Omega was used for alignment and phylogenetic tree construction. IadD from *V. paradoxus* CL014 forms a distinct subclade within the Rieske non-heme oxygenase group, with comparison to HDAO and FMO families included to contextualize functionally diverse indole oxygenases.

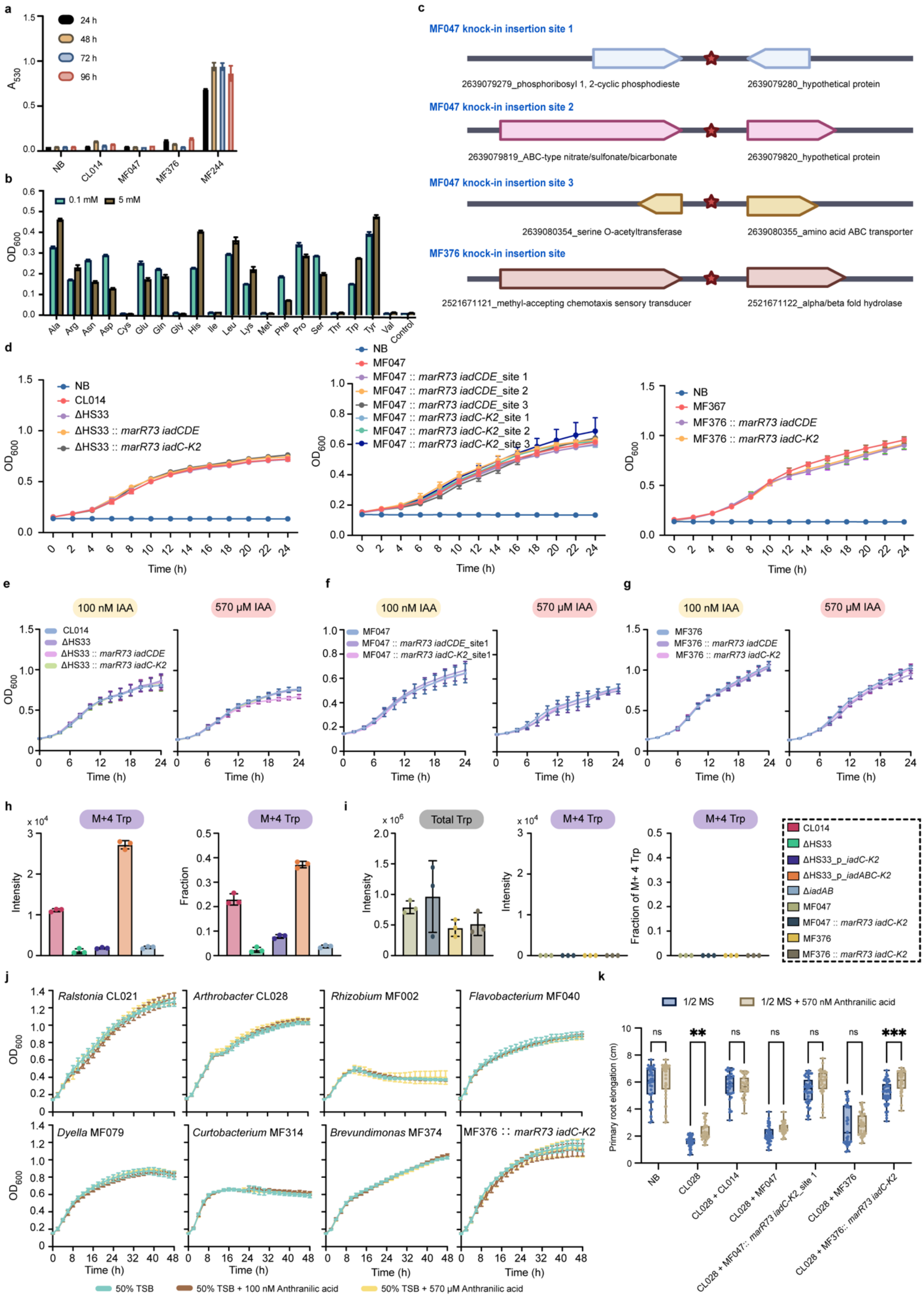
a
24 h
48 h
72 h
96 h
A530
NB
CL014
MF047
MF376
MF244
b
0.1 mM
5 mM
OD600
Ala Arg Asn Asp Cys Glu Gln Gly His Ile Leu Lys Met Phe Pro Ser Thr Trp Tyr Val Control
c
MF047 knock-in insertion site 1
2639079279_phosphoribosyl 1, 2-cyclic phosphodieste
2639079280_hypothetical protein
MF047 knock-in insertion site 2
2639079819_ABC-type nitrate/sulfonate/bicarbonate
2639079820_hypothetical protein
MF047 knock-in insertion site 3
2639080354_serine O-acetyltransferase
2639080355_amino acid ABC transporter
MF376 knock-in insertion site
2521671121_methyl-accepting chemotaxis sensory transducer
2521671122_alpha/beta fold hydrolase
d
NB
CL014
ΔHS33
ΔHS33 :: marR73 iadCDE
ΔHS33 :: marR73 iadC-K2
MF047
MF047 :: marR73 iadCDE_site 1
MF047 :: marR73 iadCDE_site 2
MF047 :: marR73 iadCDE_site 3
MF047 :: marR73 iadC-K2_site 1
MF047 :: marR73 iadC-K2_site 2
MF047 :: marR73 iadC-K2_site 3
MF367
MF376 :: marR73 iadCDE
MF376 :: marR73 iadC-K2
OD600
Time (h)
e
f
g
100 nM IAA
570 μM IAA
MF047 :: marR73 iadCDE_site1
MF047 :: marR73 iadC-K2_site1
MF376
h
i
M+4 Trp
Total Trp
Intensity
Fraction
Fraction of M+ 4 Trp
CL014
ΔHS33
ΔHS33_p_iadC-K2
ΔHS33_p_iadABC-K2
ΔiadAB
MF047
MF047 :: marR73 iadC-K2
MF376
MF376 :: marR73 iadC-K2
j
Ralstonia CL021
Arthrobacter CL028
Rhizobium MF002
Flavobacterium MF040
Dyella MF079
Curtobacterium MF314
Brevundimonas MF374
MF376 :: marR73 iadC-K2
50% TSB
50% TSB + 100 nM Anthranilic acid
50% TSB + 570 μM Anthranilic acid
k
1/2 MS
1/2 MS + 570 nM Anthranilic acid
Primary root elongation (cm)
ns
**
***
NB
CL028
CL028 + CL014
CL028 + MF047
CL028 + MF047:: marR73 iadC-K2_site 1
CL028 + MF376
CL028 + MF376:: marR73 iadC-K2

**Extended Data Fig. 6 | Engineering and characterization of *iad* knock-in strains. a**, IAA production quantified by Salkowski assay in strains grown in M9 medium supplemented with glucose (5 g $L^{-1}$) and tryptophan (5 mM). *Agrobacterium* MF224 served as a positive control. **b**, Amino acid requirements of MF047 assessed by growth in M9 medium supplemented with glucose (5 g $L^{-1}$) and individual amino acids at 0.1 or 5 mM. Data are mean ± s.d. (*n* = 2). **c**, Genomic insertion sites of *iad* constructs in MF047 (three loci) and MF376 (one locus), based on IMG annotations. Red stars indicate insertion sites. **d–g**, Growth curves of engineered CL014, MF047, and MF376 strains cultured in 50% TSB medium at 28 °C. Panel **d** shows growth in medium supplemented with IAA, whereas panels **e–g** show growth in medium lacking IAA. $OD_{600}$ was monitored over time; data are mean ± s.d. (*n* = 3-6). **h**,**i**, Isotope-tracing analysis of tryptophan production from anthranilic acid in CL014 (**h**) and engineered strains (**i**) (*n* = 3). **j**, Growth curves of representative root-associated bacterial strains cultured in 50% TSB supplemented with 100 nM or 570 μM anthranilic acid. Anthranilic acid did not noticeably affect bacterial growth under the tested conditions. Data represent mean ± s.d. (*n* = 3–4 biologically independent cultures). **k**, Primary root elongation of *Arabidopsis* seedlings grown on 1/2 MS medium with or without 570 nM anthranilic acid and inoculated with the indicated bacterial treatments. Data are from two independent experiments. Statistical significance in **k** was determined using ordinary two-way ANOVA followed by Šídák's multiple-comparisons test comparing 1/2 MS and 1/2 MS + anthranilic acid conditions within each bacterial treatment. A significant interaction between bacterial treatment and anthranilic acid treatment was detected (*P* = 0.0355). Asterisks above box plots indicate significant differences after multiple-comparisons correction (ns, not significant; ***P* < 0.01; ****P* < 0.001). Sample sizes in **k**: *n* = 39, 37, 47, 33, 31, 30, 35, 31, 39, 31, 37, 32, 45, 33.

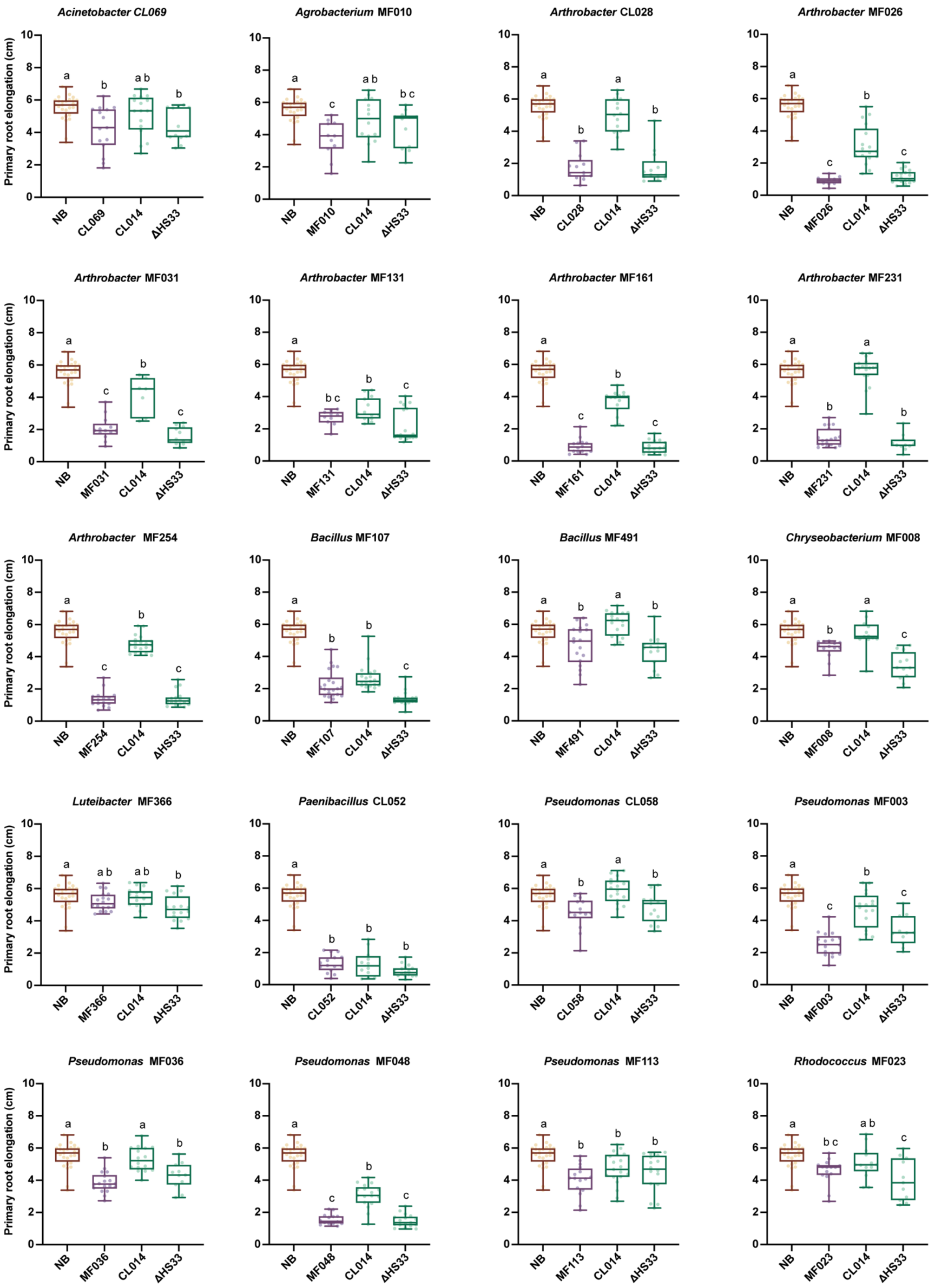
Primary root elongation (cm)
0 2 4 6 8 10
Acinetobacter CL069
a b a b b
NB CL069 CL014 ΔHS33
Agrobacterium MF010
a c a b b c
NB MF010 CL014 ΔHS33
Arthrobacter CL028
a b a b
NB CL028 CL014 ΔHS33
Arthrobacter MF026
a c b c
NB MF026 CL014 ΔHS33
Arthrobacter MF031
a c b c
NB MF031 CL014 ΔHS33
Arthrobacter MF131
a b c b c
NB MF131 CL014 ΔHS33
Arthrobacter MF161
a c b c
NB MF161 CL014 ΔHS33
Arthrobacter MF231
a b a b
NB MF231 CL014 ΔHS33
Arthrobacter MF254
a c b c
NB MF254 CL014 ΔHS33
Bacillus MF107
a b b c
NB MF107 CL014 ΔHS33
Bacillus MF491
a b a b
NB MF491 CL014 ΔHS33
Chryseobacterium MF008
a b a c
NB MF008 CL014 ΔHS33
Luteibacter MF366
a a b a b b
NB MF366 CL014 ΔHS33
Paenibacillus CL052
a b b b
NB CL052 CL014 ΔHS33
Pseudomonas CL058
a b a b
NB CL058 CL014 ΔHS33
Pseudomonas MF003
a c b c
NB MF003 CL014 ΔHS33
Pseudomonas MF036
a b a b
NB MF036 CL014 ΔHS33
Pseudomonas MF048
a c b c
NB MF048 CL014 ΔHS33
Pseudomonas MF113
a b b b
NB MF113 CL014 ΔHS33
Rhodococcus MF023
a b c a b c
NB MF023 CL014 ΔHS33

**Extended Data Fig. 7 | Primary root length of *Arabidopsis* seedlings inoculated with previously identified RGI-inducing strains[8] either alone (self) or co-inoculated with *V. paradoxus* CL014 wild type or the *iad*-deficient mutant ΔHS33, to assess *iad*-dependent reversal of RGI.** Statistical significance was determined by one-way ANOVA with Tukey's post hoc test ($P < 0.05$); groups that do not share the same letter are significantly different. "NB" denotes the no-bacteria control. Sample sizes (left to right, top to bottom): $n$ = 28, 15, 16, 12, 28, 11, 14, 12, 28, 13, 14, 13, 28, 11, 17, 18, 28, 13, 7, 12, 28, 10, 14, 18, 28, 14, 15, 17, 28, 19, 18, 11, 28, 16, 20, 18, 28, 22, 19, 16, 28, 18, 18, 15, 28, 13, 12, 12, 28, 18, 16, 16, 28, 15, 12, 17, 28, 14, 18, 13, 28, 17, 16, 11, 28, 18, 16, 11, 28, 14, 17, 17, 28, 16, 19, 16, 28, 15, 13, 11. Box plots show the median (center line), interquartile range (box), and whiskers extending to 1.5 × the interquartile range.

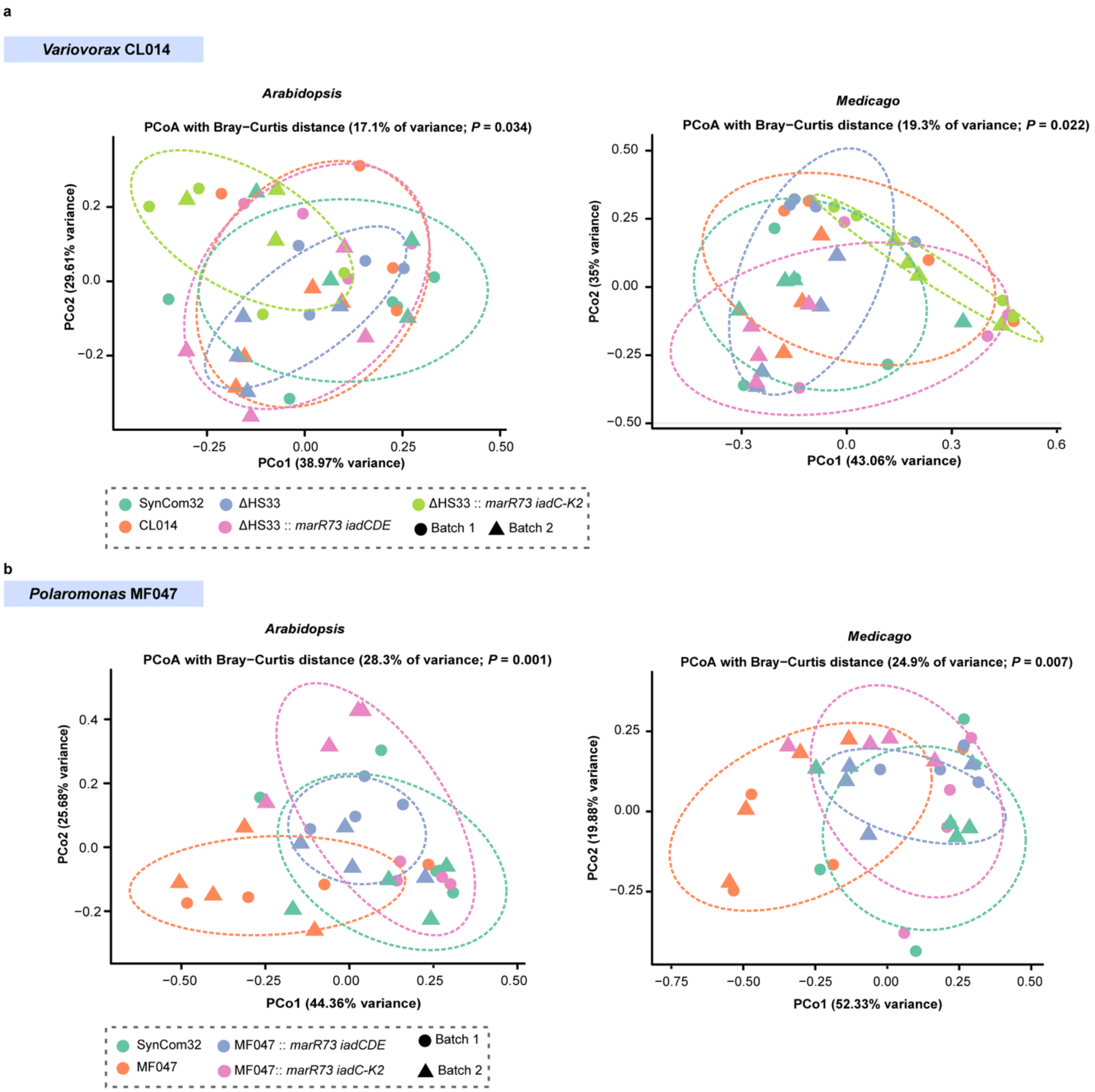


**Extended Data Fig. 8 | *V. paradoxus* CL014, *Polaromonas* MF047, and their engineered strains have modest effects on root microbiome composition.** Unconstrained principal coordinate analysis (PCoA) of Bray–Curtis dissimilarity showing root microbiome composition in *Arabidopsis* and *Medicago* treated with SynCom32 alone or co-inoculated with engineered strains of *V. paradoxus* CL014 (**a**) and *Polaromonas* MF047 (**b**). Ellipses indicate 68% confidence intervals for each treatment group. Statistical significance was evaluated using PERMANOVA (Adonis2).

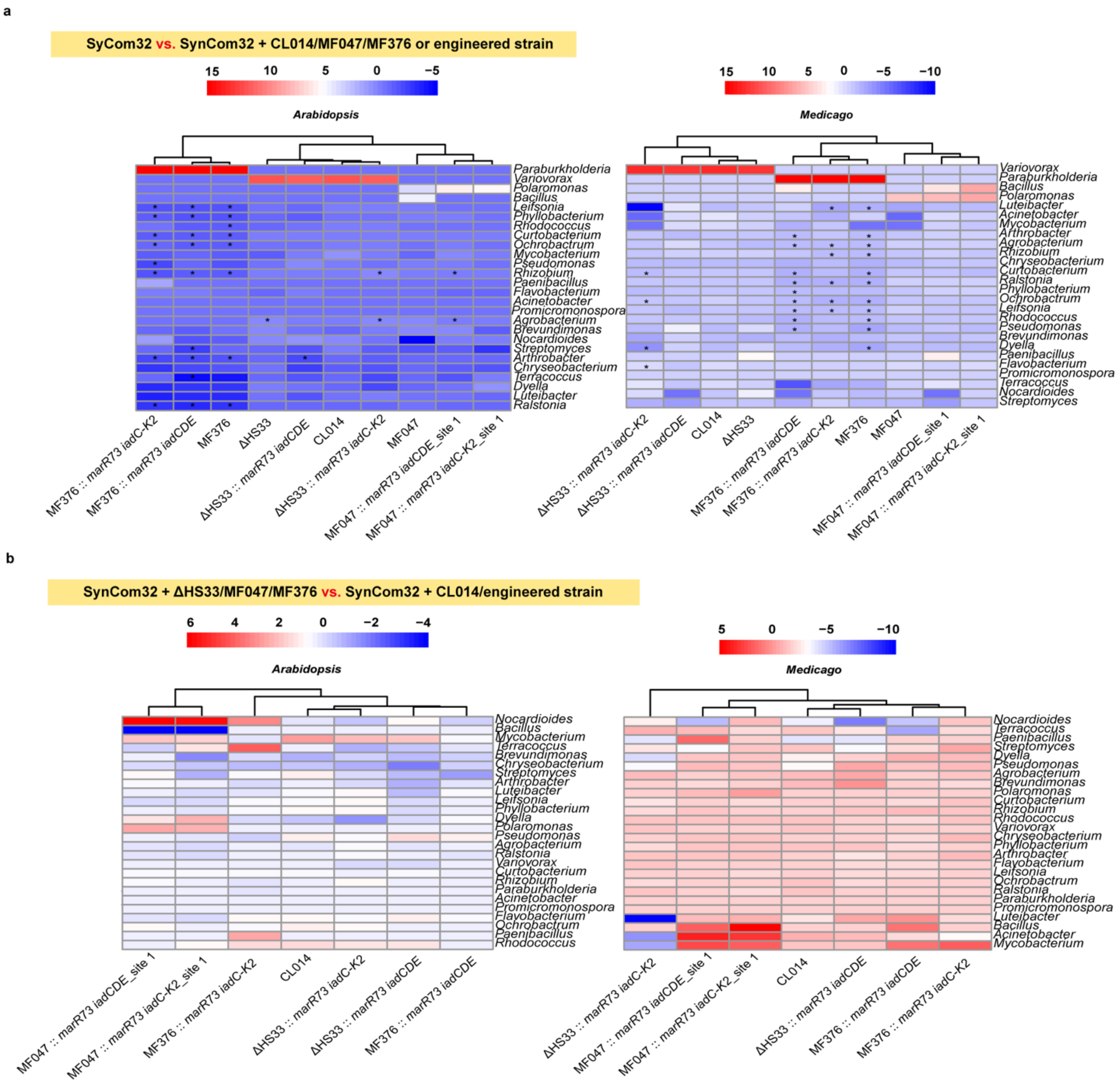


**Extended Data Fig. 9 | Log$_2$ fold-change analysis of genus-level shifts in *Arabidopsis* and *Medicago* root microbiomes.** Heatmaps show log$_2$ fold-change in bacterial genus abundance in root microbiomes treated with SynCom32 alone or co-inoculated with *V. paradoxus* CL014, *Polaromonas* MF047, *Paraburkholderia* MF376, or their engineered strains. **a**, Relative abundance changes compared to SynCom32 alone. **b**, Comparisons for each engineered strain group were made against different controls: SynCom32 + ΔHS33 for *V. paradoxus* CL014-related strains, SynCom32 + *Polaromonas* MF047 for *Polaromonas* MF047-related strains, and SynCom32 + *Paraburkholderia* MF376 for *Paraburkholderia* MF376-related strains. Pairwise comparisons were performed using Mann-Whitney U tests with FDR correction. Asterisks indicate significant differences (FDR-adjusted $P < 0.05$).

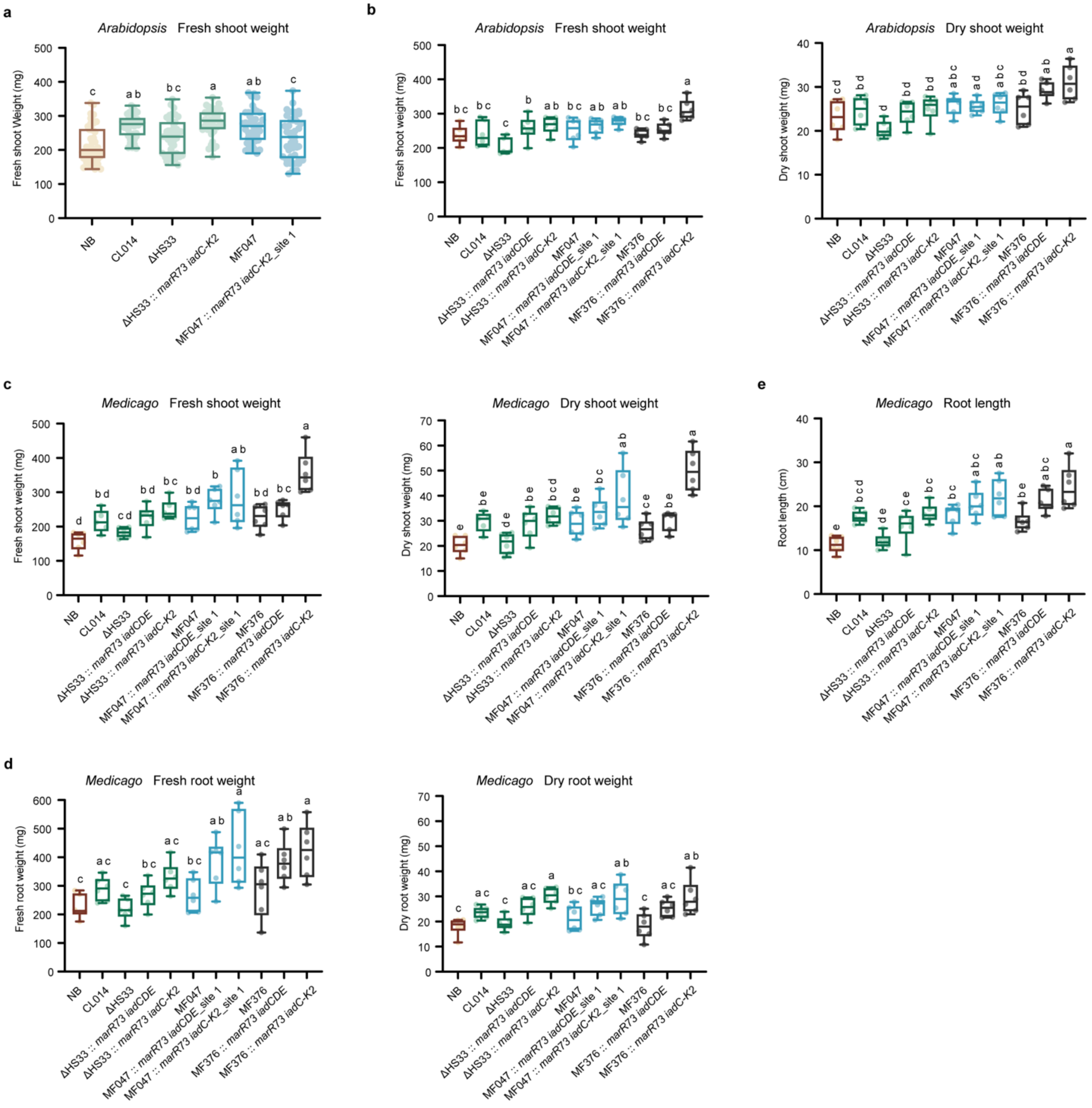


**Extended Data Fig. 10 | *Paraburkholderia* MF376 carrying the complete *iad* pathway promotes *Arabidopsis* and *Medicago* growth in natural soil after 33 days. a,b,** *Arabidopsis*; **c–e**, *Medicago*. In **a**, *Arabidopsis* shoot fresh weight data were pooled from four independent experiments, including plants grown in large pots with 3–4 plants per pot and individually grown plants from the experiment shown in **b**. In **b**, *Arabidopsis* plants were grown individually in small pots to facilitate root recovery, but intact roots could not be reliably obtained; therefore, only shoot traits were quantified. For *Medicago*, plants were grown individually and shoot, and root traits were quantified (**c–e**). Letters above box plots indicate statistically significant differences determined by one-way ANOVA with Tukey's post hoc test on the underlying replicate values ($P < 0.05$). Sample sizes: **a**, $n$ = 42, 40, 41, 38, 43, 41; Data were obtained from four independent experiments. **b–e**, each treatment contained 5–6 samples. Data were obtained from two independent experiments.
